# EEG Microstate Correlates of Emotion Dynamics and Stimulation Content during Video Watching

**DOI:** 10.1101/2021.10.09.463281

**Authors:** Wanrou Hu, Zhiguo Zhang, Huilin Zhao, Li Zhang, Linling Li, Gan Huang, Zhen Liang

## Abstract

EEG microstates have been widely adopted to understand the complex and dynamic-changing process in dynamic brain systems, but how microstates are temporally modulated by emotion dynamics is still unclear. An investigation of EEG microstates under video-evoking emotion dynamics modulation would provide a novel insight into the understanding of temporal dynamics of functional brain networks. In the present study, we postulate that emotional states dynamically modulate the microstate patterns, and perform an in-depth investigation between EEG microstates and emotion dynamics under a video-watching task. By mapping from subjective-experienced emotion states and objective-presented stimulation content to EEG microstates, we gauge the comprehensive associations among microstates, emotions, and multimedia stimulation. The results show that emotion dynamics could be well revealed by four EEG microstates (MS1, MS2, MS3, and MS4), where MS3 and MS4 are found to be highly correlated to different emotion states (emotion task effect and level effect) and the affective information involved in the multimedia content (visual and audio). In this work, we reveal the microstate patterns related to emotion dynamics from sensory and stimulation dimensions, which deepens the understanding of the neural representation under emotion dynamics modulation and will be beneficial for the future study of brain dynamic systems.

## 1 Introduction

Electroencephalography (EEG) microstate analysis is an effective computational method that provides perspectives into EEG spatial-temporal dynamics (Koenig et al. 2002; Khanna et al. 2015; Michel and Koenig 2018). Microstates, as a topographical information representation of the electrical potentials over multichannel EEG signals, offer a powerful approach to reveal the spatial-temporal dynamical system of whole-brain spontaneous activities. Recent findings show EEG microstates have close electrophysiological relations with global functional brain networks observed from functional magnetic resonance imaging (fMRI). For the commonly recognized four canonical microstates (termed as **MS1, MS2, MS3, MS4** below), the existing studies show the four microstates are associated with **auditory, visual, default mode, and dorsal attention networks** (Koenig et al. 2002; Khanna et al. 2015; Michel and Koenig 2018).

Microstates reflect momentary brain activities with high temporal resolution. The changes of microstates do not happen randomly, and the transitions between microstates could be interpreted as a sequential activation of different global functional brain networks. Current microstate studies have found microstates can directly characterize the qualitative and quantitative aspects of perception and cognition processing (Britz et al. 2010; Musso et al. 2010; Yuan et al. 2012). For example, Seitzman et al. (2017) discovered that cognitive task manipulation yielded a significant increase in MS4 but a decrease in MS3. Besides, the visual-stimulated tasks were mainly related to MS2 activities. In Milz et al.’s work (2016), MS1 and MS2 activities were found to be significantly related to the visual and verbal processing in the brain, and MS4 mainly responded during goal-directed perceptual task processing. These results suggest that a representation of the quasi-stable and consistent set of patterns could be established as reliable neurophysiological biomarkers for brain activity study under various physiological and psychological states.

Till now, the given physiological and cognitive significance of EEG microstates has not been fully explored. The emotional state could be probably an important factor modulating the representation patterns of microstates and affecting the microstate transitions. In Gianotti et al.’s (2008) emotion-evoked event-related potential (ERP) work, it was found the temporally microstate dynamic patterns on valence and arousal emotion dimensions were different. Valence-related microstate was first detected around 118 ms in emotional word experiments and around 142 ms in emotional picture experiments; arousal-related microstate was found early at 266 ms in word experiments and 302 ms in picture experiments. The results observed an earlier effect of valence states on EEG microstate activities than the arousal and demonstrated the effectiveness of EEG microstate analysis in emotion study. Besides, Shen et al. (2020) discovered the microstate syntax was discriminant for emotion understanding, where the transition probability from one microstate to another was correlated with emotion ratings in terms of valence and arousal. Here, the public microstate templates were adopted for microstate detection and possible microstate parameters such as duration, occurrence, coverage, and transition probabilities were all considered. These works show there exists a link between the alternations of microstate presentations and emotions. However, existing studies concerning the associations between microstates and emotions are still quite limited. Much less is known about how time-series microstate representations dynamically alter with the subjective-experienced emotions. An in-depth neurophysiological investigation is required to lead to a more complete, accurate, and reliable deciphering of the temporally modulation effect of emotions on microstates.

On the other hand, video is widely used as experimental stimulation in neurophysiological experiments for emotion induction, which contains rich audiovisual information and offers realistic and vivid scenarios for evoking various human emotions in a laboratory environment (Koelstra et al. 2012; Zheng and Lu 2015; Katsigiannis and Ramzan 2018). During video watching, EEG signals related to the video-evoking emotions are simultaneously recorded. Here, to wider our knowledge of the emotion-related EEG dynamics in terms of microstates and examine whether the sentiment information involved in the multimedia stimulation could also modulate the changing microstates, an investigation of the association between the evoked emotional dynamic EEG microstate activities and video content is carried out.

Overall, to better understand emotion-induced microstates, the present work investigates the involved potential emotion relevance in EEG microstate representations in terms of **subjective-experienced emotion state analysis (Study 1)** and **objective-presented stimulation effect analysis (Study 2)**. The main contribution of this work is to understand how EEG microstates are temporally modulated by emotions. The underlying relationships among microstate dynamics, subjective-experienced emotion state, and objective-presented multimedia stimulation content are investigated. Through mapping from subjective-experienced emotion states to neurophysiological signals and from objective-presented stimulation content to real-perceived emotion processing in the brain, this work offers a deeper insight into which characteristics of microstates are relevant to potential emotion changes.

## 2 Methods

### 2.1 Overview

In light of the sensitivity of microstate parameters to transient changes in brain states, the relations among EEG microstates, emotion states, and stimulation content are explored, and a clarification of how microstates related to the internal (emotions) and external (stimulation) changes will be conducted (as shown in Fig. 1). For the subjective-experienced emotion state analysis (Study 1), an examination of the changes in microstates related to evoked emotion states will be implemented. In line with the complex characteristics of emotions, we will manifest a comprehensive study on emotion dynamics from the representation patterns of EEG microstates under different emotional states (**task effect**), emotional levels (**level effect**), and during processes of emotion induction (**evoking dynamics**). For the objective-presented stimulation effect analysis (Study 2), the temporal associations between EEG microstate activities and multimedia stimulation will be estimated to an alternative understanding of how emotion perceives in the brain. The evoking effect of emotional multimedia will be separately analyzed on visual and audio content. In summary, two studies will be conducted as follows.

- **Study 1: subjective-experienced emotion state analysis** (**Section 2.4**). EEG microstate dynamics will be first characterized to describe spatial-temporal changes of EEG activities under various emotion states during video watching. Subsequently, the activity patterns of EEG microstates will be analyzed to characterize emotion-related neural dynamics from three perspectives. (1) **Task effect** (Section 2.4.1): to quantify the differences of EEG microstate activities between pre-stimulus and post-stimulus stages. (2) **Level effect** (Section 2.4.2): to measure the differences of EEG microstate activities between low-level and high-level groups on different emotion dimensions. (3) **Evoking dynamics** (Section 2.4.3): to evaluate the temporal dynamics of emotion processing in the brain, in which the moment-to-moment changes of EEG microstate activities during video watching will be examined.
- **Study 2: objective-presented stimulation effect analysis** (**Section 2.5**). The multimedia content for emotion induction will be described by the low-level attribute features and high-level semantic features of **visual content** (Section 2.5.1) and **audio content** (Section 2.5.2). The corresponding associations between visual/audio content and emotion-related EEG microstate activities will be measured. The **timing effect** (Section 2.5.3) of visual and audio content on time-series microstate representation will be separately studied, and the temporal correlations between the dynamic EEG microstate activities and the changing visual/audio content will be examined.

**Fig. 1.**
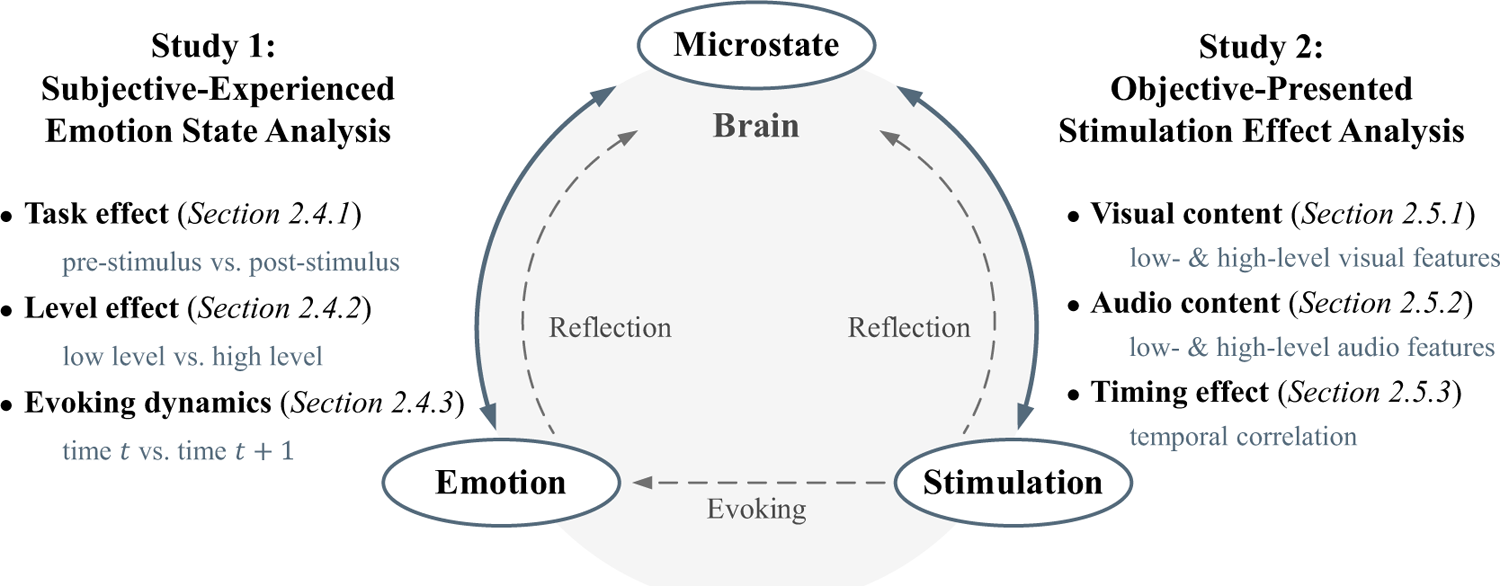
The investigation framework of EEG microstates under modulation of video-evoked emotion dynamics. In Study 1, based on EEG microstate analysis, the relationships between subjective-experienced emotion states and dynamic brain activities are characterized. The emotion-related EEG dynamics are described by the representation pattern of EEG microstate activities under the exploration of task effect, level effect, and evoking dynamics. In Study 2, the stimulation effect of objective-presented multimedia content on EEG microstate activity patterns is analyzed. The time effect of emotional stimulation is to evaluate temporal associations between EEG microstate activities and presented visual/audio content in a time-shifting manner.

### 2.2 EEG data and preprocessing

A well-known DEAP database (a database for emotion analysis using physiological signals) (Koelstra et al. 2012) with 32 subjects’ EEG recordings during video watching is used for emotion dynamics investigation. In this database, 40 emotional music videos (corresponding to 40 trials below) with a fixed length of 60 s were randomly presented for emotion induction. Simultaneously, 32-electrode EEG signals were recorded at a sampling rate of 512 Hz. Throughout the entire emotion-evoking experiment, all the EEG data were collected in an eye-open condition. To investigate the emotion-related brain dynamics from the recorded EEG data, we first perform a standard EEG preprocessing procedure (including filtering, common average re-reference, and independent component analysis) for noise removal and signal quality enhancement. Then, the preprocessed EEG data are divided into **pre-stimulus** (3 s), **video-stimulated** (60 s), and **post-stimulus** (3 s) segments for further analysis. More details about the DEAP database and EEG preprocessing procedure are provided in Appendix I of the Supplementary Materials.

### 2.3 EEG microstate analysis

In this section, we will first introduce a standard EEG microstate analysis. In the EEG microstate analysis, microstate detection is a fundamental and crucial part, which will directly influence the validity and reliability of the analysis performance. However, the current microstate detection methods mainly focus on the resting-state EEG and may fail to be adaptive enough for the task-state. In this work, we introduce a sequential microstate clustering analysis for efficient and representative EEG microstate template detection.

A standardized EEG microstate analysis is implemented as shown in Fig. 2, including candidate topography extraction, microstate detection, back-fitting, and feature extraction (Pascual-Marqui et al. 1995). EEG microstate analysis starts with a bottom-up extraction of EEG microstate templets from the spontaneous EEG signals (Fig. 2 (b) and (c)). Then, a top-down process termed back-fitting (Fig. 2 (d)) is conducted to re-represent EEG data into a series of dynamic microstate sequences. Based on the criterion of global map dissimilarity (GMD) (Murray et al. 2008), each EEG sample point is assigned to one microstate with high spatial similarity (Lehmann et al. 2005; Zanesco et al. 2020). Then, a temporal smoothing process (Poulsen et al. 2018) is adopted to reject the noisy time segments, and the interrupted microstate segments shorter than 30 ms would be re-assigned to another microstate based on GMD calculations. Finally, the corresponding microstate features are extracted to quantify the dynamic changes of EEG microstate activities (Fig. 2 (e)) and represent the spatial-temporal oscillations of brain activities during video watching.

**Fig. 2.**
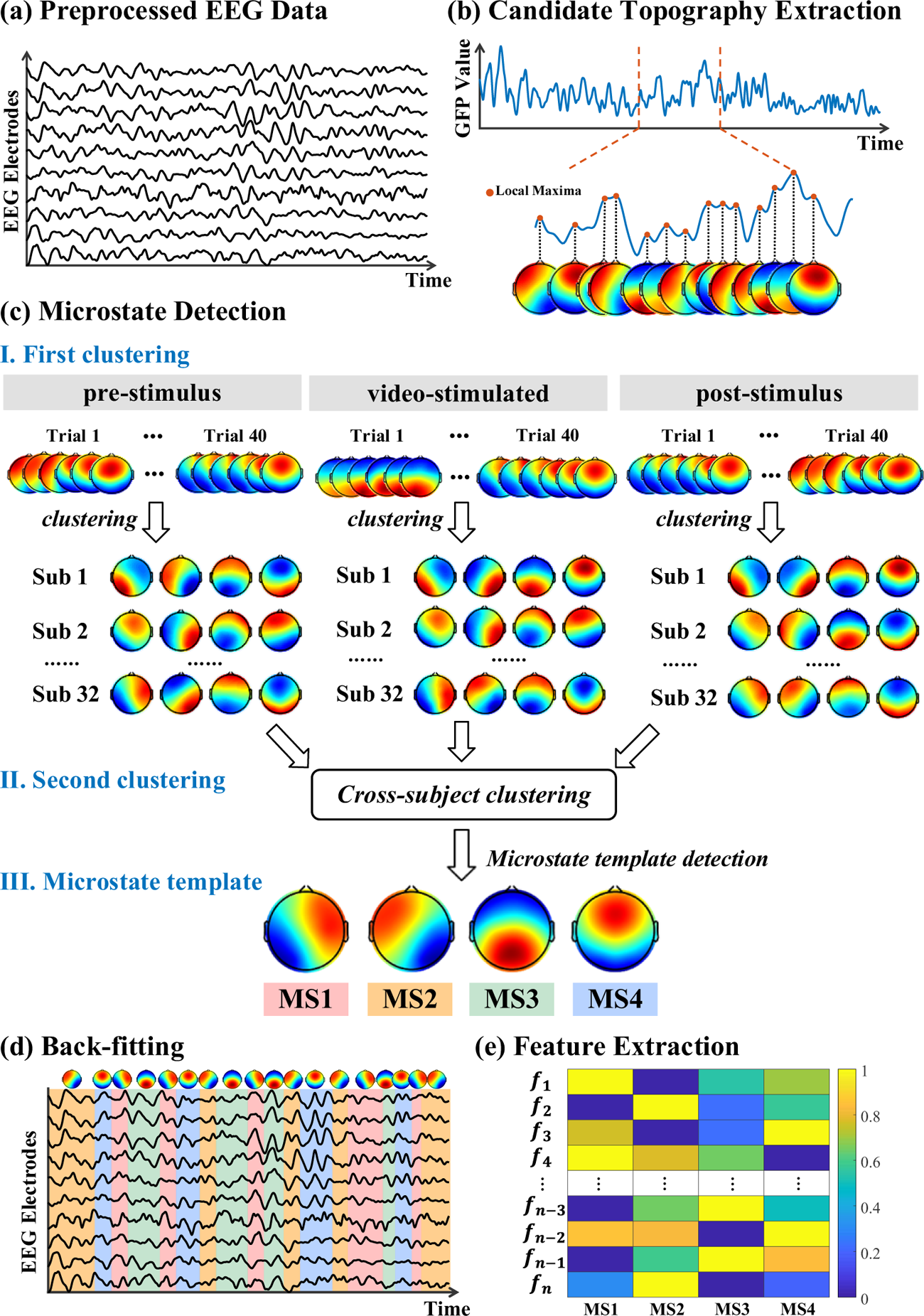
A standard procedure of EEG microstate analysis. Based on (a) the preprocessed EEG data, (b) candidate topographies with high signal-to-noise ratios are extracted from the local maxima of the GFP curve. (c) A sequential microstate clustering is conducted for representative and reliable EEG microstate detection, where the first clustering is performed at a within-subject level and the second clustering is conducted at a cross-subject level. (d) The final detected EEG microstate templates are then fitted back into the preprocessed EEG data by assigning each time point to one predominant microstate. After EEG microstate back-fitting, the original EEG time series are re-represented into EEG microstate sequences covering whole-brain spontaneous spatial-temporal activities. (e) A series of microstate features are calculated for quantitative measurement, including duration, occurrence, coverage, and transition probability, named as {*f_1_,…,f_k_,…,f_n_*}. Here, *f_k_* refers to the kth extracted microstate feature and *n* is the total feature number.

To improve the representative microstate template detection for emotion-related EEG dynamics analysis, a sequential microstate detection with two-step spatial clustering is introduced (Fig. 2 (c)). The first clustering is implemented at a within-subject level to extract subject-representative microstate topographies from both eye-open resting-state EEG data (pre-stimulus and post-stimulus stages) and task-state EEG data (video-stimulated stage). Here, based on the recorded 40 trials of EEG data from every single subject, the candidate topographies under three experimental stages are separately extracted from the local maximum of global field power (GFP) (Lehmann and Skrandies 1980; Skrandies 1989) and then output into a modified k-means clustering algorithm (Pascual-Marqui et al. 1995) with a cluster number *c* ranging from 2 to 8 and an iteration number *I* of 1000. Within the iteration of spatial clustering, the cluster centroids with high global explained variance (GEV) and low cross-validation (CV) criterion values are identified and extracted as subject-representative microstate topographies. In other words, the subjective-representative microstate topographies are identified in a data-driven manner with optimal microstate numbers and topographical shapes (as shown in Appendix II of the Supplementary Materials). Next, following the similar parameter settings of the first-step clustering, the second-step clustering is conducted for final EEG microstate template detection. Through a clustering of all the identified subject-representative microstate topographies across 32 subjects and three different experimental stages, the final detected EEG microstate templates could be established as reliable neurophysiological patterns to represent the EEG changes under emotion induction. The EEG topographies with high spatial similarity are identified as the final EEG microstates for emotion-related neural dynamics analysis. The obtained EEG microstate templates (named MS1, MS2, MS3, and MS4) show a low variance of residual noise (Pascual-Marqui et al. 1995; Murray et al. 2008) and good tolerance for individual differences (Michel and Koenig 2018; D’Croz-Baron et al. 2021) in emotion-evoked EEG dynamics analysis across 32 subjects and 40 trials. After back-fitting using the detected EEG microstates, a series of EEG microstate time sequences are obtained. Four commonly used EEG microstate features are extracted below for further investigation of emotion-related neural mechanisms.

- **Duration:** the average time span that a specific microstate remains dominant, which can reflect the stability of the underlying neural configuration during emotion induction (Khanna et al. 2015).
- **Occurrence:** the times of presentation per second that a specific EEG microstate remains dominant, which indicates the representation tendency of the underlying neural activation (Koenig et al. 2002).
- **Coverage:** the ratio of the period that a specific microstate keeps dominant to the total recording time (Seitzman et al. 2017).
- **Transition probability (TP)**: the transition percentage between any two EEG microstates, which estimates the sequential activation tendency of scalp electric potentials on a millisecond time scale (Koenig et al. 2005; Khanna et al. 2015).

### 2.4 Study 1: subjective-experienced emotion state analysis

In Study 1, we will investigate the representation differences of emotion-evoked EEG microstate activities from the perspectives of (1) task effect, (2) level effect, and (3) evoking dynamics.

#### 2.4.1 Task effect

The emotion task effect on EEG microstate activities is measured as the pairwise statistical differences between pre-stimulus and post-stimulus stages (Fig. 3 (a)). First, for each subject, the representation differences of EEG microstate activities before and after emotion-evoking tasks are measured as *d^i^_j_* (*i* ∈ [1,32] is the subject number and *j* ∈ [1,40] is the trial number). The pairwise differences between pre-stimulus and post-stimulus stages are first measured. Specifically, the differences in terms of each microstate feature extracted at the pre-stimulus and post-stimulus stages from the same trial are calculated. The pairwise differences across 40 trials for each subject are then output for t-statistic calculation. A t-value, *t_i_* (*i* ∈ [1,32]), is calculated as the statistical measurement of subject-specific representation differences in one type of microstate feature. Thus, for one microstate feature, a list of t-values from 32 subjects is obtained. Second, to evaluate the emotion-evoking task effect from a cross-subject perspective, a one-sample t-test is implemented on the obtained list of t-values for one microstate feature and a p-value is measured. Third, to minimize the influence of type I errors, all the obtained p-values are corrected for multiple comparisons using the false discovery rate (FDR) with a significance level of 5% (*p* < 0.05).

**Fig. 3.**
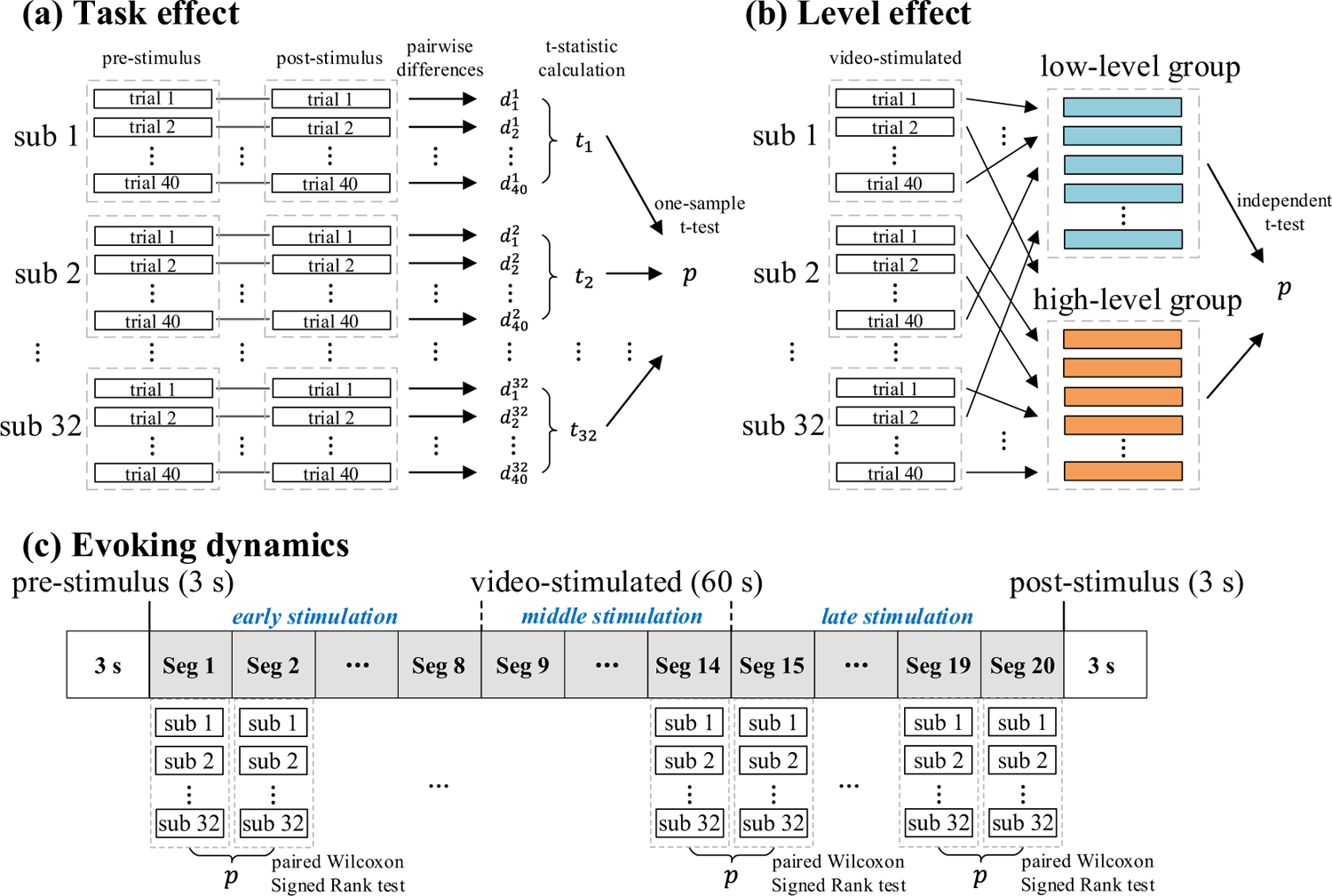
The statistical analysis processes of emotion dynamics analysis in terms of (a) task effect, (b) level effect, and (c) evoking dynamics.

#### 2.4.2 Level effect

The emotion level effect characterizes the EEG microstate differences at different levels (high or low) on emotion dimensions (valence and arousal). Generally, the level differences in valence reflect the category of emotions ranging from negative to positive states and the level differences in arousal reflect the extent of evoked emotions ranging from boring to exciting states (Morris 1995; Kensinger 2004; Alarcão and Fonseca 2019). The level effect on every emotion dimension offers alternative inspects into the neural mechanism of emotion perception. For each emotion dimension, we first divide the emotion-evoked EEG data into low- and high-level emotional groups according to the self-assessment ratings from the 32 subjects. Here, the threshold for level grouping is defined by a self-adaptive threshold reassignment method (Yin et al. 2017) that is presented in detail in Appendix III of the Supplementary Materials. Second, the emotion level effect is measured as the representation difference between low- and high-level groups in terms of each microstate feature (Fig. 3 (b)). As the emotion dynamics on different emotion dimensions are independent, the evaluation of the emotion level effect is separately measured on valence and arousal dimensions. Specifically, based on the distribution estimation of a Lilliefors test, an independent t-test (for normally distributed groups) or Wilcoxon Rank Sum test (for non-normally distributed groups) is conducted for statistical analysis on each microstate feature and the inter-group differences across trials and subjects are measured. The obtained p-values reveal the statistical evaluation of emotion level effect on EEG microstate activities.

#### 2.4.3 Evoking dynamics

In emotion-evoking experiments, continuous presentation of multimedia stimulation dynamically influences brain activities during stimulation perception and processing (Zheng and Lu 2015). An investigation of temporal variations in multimedia-evoked EEG activities helps to understand the dynamic characteristics of emotion processing in the brain. In this study, we measure the temporal dynamics of EEG activities under each video of emotion-evoking tasks in terms of each microstate feature as follows (Fig. 3 (c)). **(1) Data segmentation.** The video-stimulated EEG data at one trial is divided into a number of short segments with a fixed length of 3 s. Total 20 segments are obtained (video length 60 s / segment length 3 s = 20 segments). There is no overlap between any two adjacent segments. To highlight the temporal variation of emotion-related EEG dynamics, for clarity, the first eight segments are named as **early stimulation stage (1∼24 s)**, the following six segments are marked as **middle stimulation stage (25∼42 s)**, and the last six segments are assigned to **late stimulation stage (43∼60 s)**. **(2) Segment-based feature extraction.** Four types of microstate features (duration, occurrence, coverage, and TP) are extracted from each segment. **(3) Baseline correction.** To obtain a better estimation of emotion dynamics and minimize the emotion-unrelated effect, a baseline correction is employed by normalizing the segment-based microstate features with the corresponding features extracted from the pre-stimulus stage (−3 to 0 s). **(4) Statistical measurement.** For each video, a paired Wilcoxon Signed Rank test (non-parametric statistical analysis based on the normality estimation of Lilliefors test) is conducted on any two adjacent segments across 32 subjects (stimulated under the same given video) in terms of each microstate feature, and 19 pairs (total 20 segments for each video) of segment-based statistical differences are obtained to represent the video-specific temporal characteristics of emotion processing in the brain.

### 2.5 Study 2: objective-presented stimulation effect analysis

To better understand the neural dynamics in emotion induction, the stimulation effect of objective-presented multimedia content on EEG microstate activity changes is further analyzed through a temporal correlation analysis following an analytical process shown in Fig. 4. In line with the characteristics of human sensory processing, both low-level attribute features and high-level semantic features from visual and audio content are extracted to give a fuller description of the stimulation content delivered in a given video clip. Taking advantage of the development of computer science and deep learning fields, the high-level visual and audio features are characterized by two pre-trained deep convolutional neural networks (CNNs).

**Fig. 4.**
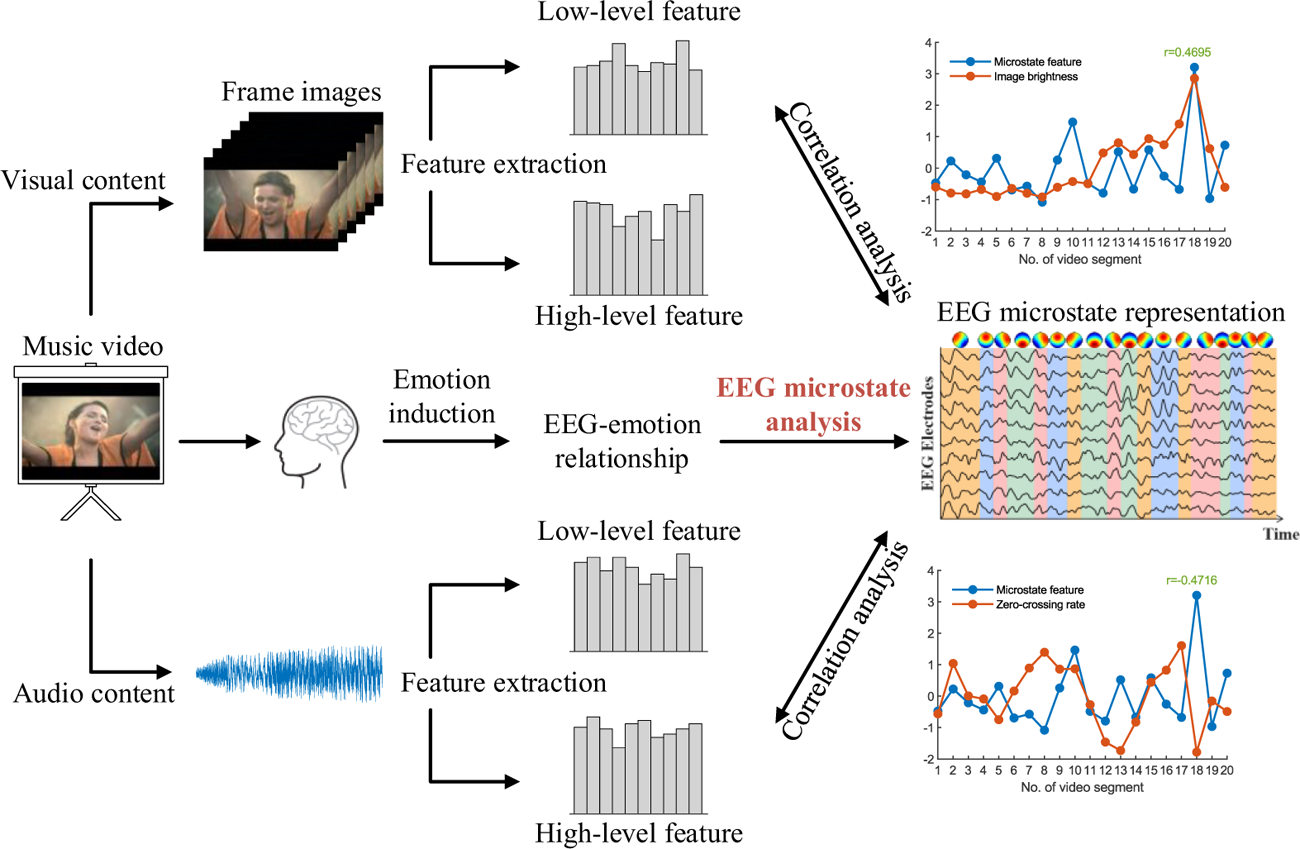
A general flowchart for temporal correlation detection between emotion-related EEG microstate activities and emotion-evoking multimedia stimulation content. In this process, how EEG microstates react to different objective-presented multimedia stimulation content is studied for emotion perceptual mechanism investigation.

#### 2.5.1 Visual content

Image brightness is a fundamental visual characteristic for visual perceptual processing (Itti et al. 1998) with a well-validated role in emotion induction (Lakens et al. 2013). In this study, the brightness of video frames is extracted as the low-level visual feature for attribute representation. Frame-based visual brightness is first extracted and then converted to a segment-based visual feature through an average calculation. The changes of segment-based visual features characterize the temporal variations of low-level visual content in multimedia stimulation, which provides the evoking clues for emotion dynamics analysis.

A pre-trained VGG16 model (Simonyan and Zisserman 2014), a well-recognized CNN model with 13 convolutional layers, is utilized for high-level visual feature extraction. The VGG16 model was trained by millions of labeled categorial images from the ImageNet database (Deng et al. 2009), which is capable of extracting discriminative features for visual representation and achieves outstanding performance in image classification. Each convolutional layer of the VGG16 model functions as an automated feature extractor. With an increase of convolutional layers, from the shallow to the deep layers, more semantic related visual information could be successfully characterized. To extract high-level visual features for emotion dynamics analysis (Fig. 5 (a)), video frames are sequentially input into VGG16, and a set of feature maps are characterized from the deep convolutional layers (termed as *conv*11, *conv*12, and *conv*13) that corresponding to the semantic information involved in the visual content. Similar to low-level visual feature extraction, frame-based visual features are obtained and then converted to segment-based visual features through an average calculation. In total, one low-level visual feature and three types of high-level visual features are extracted for characterizing the visual content of the video clip in terms of segments.

**Fig. 5.**
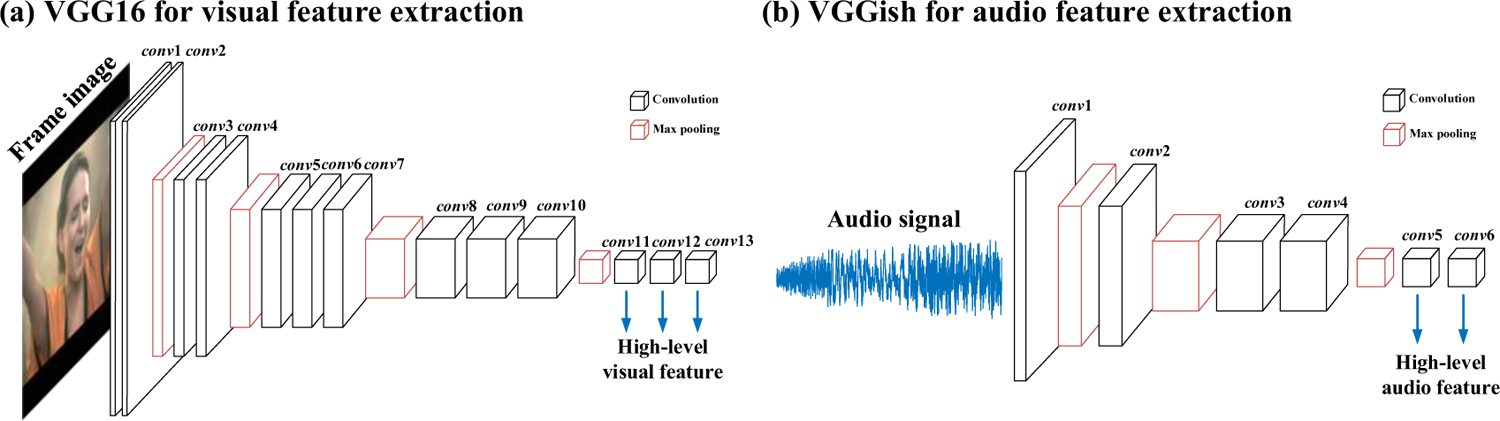
The pre-trained deep convolutional neural networks (VGG16 and VGGish) for high-level visual and audio feature extraction. For visual content, the features extracted from *conv*11 to *conv*13 in VGG16 are termed high-level visual features. For audio content, the features extracted from *conv*5 and *conv*6 in VGGish are termed high-level audio features.

#### 2.5.2 Audio content

Zero-crossing rate (ZCR) (Teixeira et al. 2012) is an inherent prosodic feature that characterizes the frequency component of the audio signal by counting the average number that the audio amplitude crosses zero within a given time interval. In this study, the ZCR feature is extracted at each segment as the low-level audio feature, and the temporal variations among the segment-based audio features quantify the audio content presented for emotion induction.

To characterize the semantic content delivered by the audio content, a pre-trained VGGish (Hershey et al. 2017) with 6 convolutional layers is adopted for high-level audio feature extraction. The VGGish model was trained by 2 million manual-labeled YouTube video soundtracks from the Google Audio Set (Gemmeke et al. 2017), and it has been proven to be an excellent audio feature extractor for learning representative deep characteristics from the original video soundtracks. For each video segment, the corresponding soundtrack signal is input into the VGGish model and the high-level audio features for sematic content representation are extracted at the 5^th^ and 6^th^ convolutional layers (termed as *conv*5 and *conv*6 in Fig. 5 (b)). In total, for each audio segment, there are one low-level audio feature and two types of high-level audio features.

#### 2.5.3 Timing effect

To interpret the dynamic process of multimedia-evoked emotion induction, the timing effect of stimulation content on EEG microstate activities is investigated. In a time-shifting manner, the time courses of visual and audio features are shifted in a preceded or succeeded direction. The temporal relationships are examined between presented emotional multimedia content and real-evoked EEG microstate activities. Here, the shifting range is given from −1 s (stimulation preceded) to 1 s (stimulation succeeded), with a step of 100 ms. In total, there are 21 time-shifting parameters. Note that no time-shifted processing is applied on EEG microstate features. For each time-shifting parameter, the temporal correlation between the shifted multimedia stimulation and EEG microstate responses is measured as follows. **(1) Temporal brain dynamics computation.** For each trial, the subject-specific time-varying EEG microstate activities are characterized by calculating the first-order differences between any two adjacent segments in terms of each microstate feature. **(2) Temporal multimedia dynamics computation.** For each multimedia stimulation, the contents are first shifted at the given time-shifting parameter. Then, the temporal changes are measured by computing the first-order differences in terms of segment-based visual or audio features. **(3) Subject-specific correlation measurement.** For each subject and each microstate feature, the temporal correlation between the computed temporal brain dynamics and the computed temporal multimedia dynamics (with a specific time-shift parameter) is measured by calculating the Pearson correlation coefficients. For each multimedia stimulus, in total 32 subject-specific correlation coefficients (corresponding to 32 subjects) are obtained for each microstate feature and for each time-shifting parameter. **(4) Video-specific correlation measurement.** For each multimedia stimulus and each microstate feature, the obtained 32 subject-specific correlation coefficients are then verified by t-statistic calculation (cross-subject measurement). The calculated t-values quantify the general changing trend of EEG microstate activities in response to the given multimedia stimulation (after time-shifting) across 32 subjects. **(5) Cross-video temporal correlation evaluation.** To explore the cross-video and cross-subject stimulation effect on the dynamic changes of each EEG microstate feature, the calculated 40 t-values (corresponding to 40 videos) in step (4) are then fed into a two-tailed one-sample t-test. The final obtained t-value characterizes the temporal stimulation effect on EEG microstate activities with a positive or negative correlation, and the corresponding p-value reveals whether the temporal stimulation effect is statistically significant. The above steps (1)-(5) are repeated until each time-shifting parameter and each microstate feature are evaluated, and the overall timing effect of multimedia stimulation on emotion-related EEG dynamics is obtained.

## 3 Results

### 3.1 Overview

In this section, we will present the observations of the changes in microstate parameters under the studies of subjective-experienced emotion state analysis (Study 1) and objective-presented stimulation effect analysis (Study 2). Two main observations of this work are summarized as follows. (1) The representation patterns of EEG microstates can reveal the subjective-experienced emotion states. Different distinctive microstate patterns are observed for different emotion dimensions. It is found that arousal changes mainly affect MS3 activities, and valence states are mainly related to MS4 activities. (2) Objective-presented multimedia stimulation content modulates the representation pattern of emotion-related EEG microstate activities. The changes of microstates perform differently for perceiving different stimulation information. Visual content mainly leads to the changes in MS4 activities, and audio content is mainly related to the changes in MS3. Overall, by examining the dynamic characteristics of brain activities during emotion induction, the associations among EEG microstates, emotion states, and stimulation content are verified.

### 3.2 EEG microstates

Based on the DEAP database, four EEG microstates are detected in a data-driven manner as presented in Section 2.3. The corresponding GEV and CV values of four identified microstate templates are 82.23% and 63.33% (as shown in Fig. 6 (a)), referring to the maximum first-order difference in a ratio of GEV to CV (Fig. 6 (b)). The detected microstate templates are presented in Fig. 6 (c), which share similar topographical configurations to the canonical microstates reported in the literature (Pascual-Marqui et al. 1995; Koenig et al. 2002; Britz et al. 2010; Michel and Koenig 2018). For consistency, we label the detected microstate templates as MS1, MS2, MS3, and MS4 according to topographical orientation. Besides, to quantify the stability of the detected microstate templates along with emotion-evoking experiments, we introduce two evaluation indexes (microstate proportion and Cronbach’s α value) for quantitative measurement of the 4 microstate templates. As reported in Table 1, MS3 and MS4 account for a major proportion across three experimental stages, compared to MS1 and MS2. The calculated proportion value of MS1 and MS2 is 37.71±0.76% and that of MS3 and MS4 is 62.30±0.76%. A larger proportion of MS3 and MS4 is in line with the finding that MS3 and MS4 activities are more related to emotion perception in the brain. The Cronbach’s α values across three experimental stages are all larger than 0.7 (0.7290±0.0239), supporting that the identified EEG microstates are temporally stable along with emotion-evoking experiments.

**Fig. 6.**
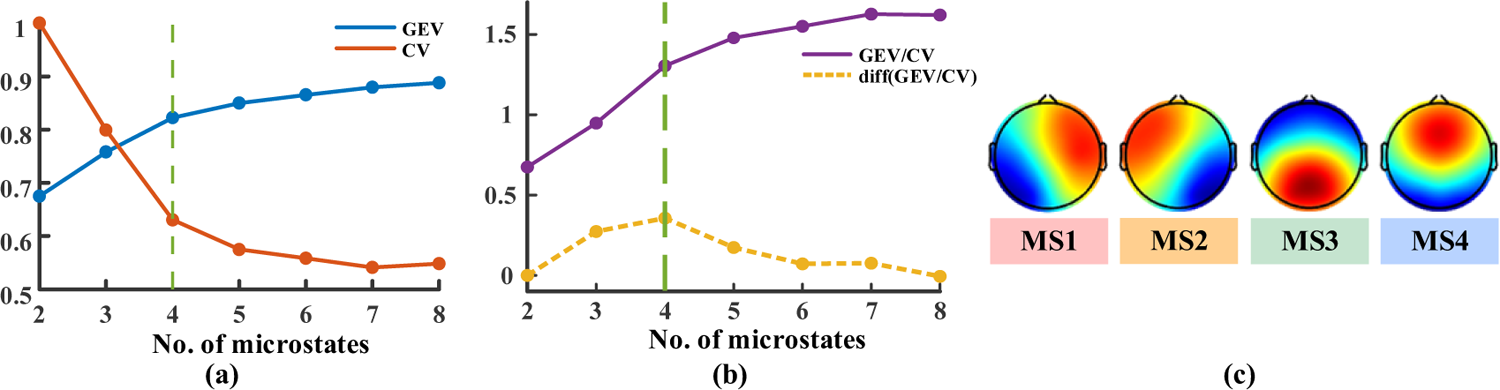
The detected EEG microstate templates for emotion-related EEG dynamics analysis. (a) Microstate detection performance with different numbers of EEG microstate classes and the calculated GEV and CV values. Here, the corresponding calculated GEV and CV values of the four identified microstate templates are 82.23% and 63.33%. For visualization, the CV values are normalized to the range of [0, 1]. (b) Under consideration of GEV and CV values in the interaction process of topographical clustering, an optimal cluster number of four is identified which refers to the maximum first-order difference in a ratio of GEV to CV. (c) Visualization on the topographical shape of the final detected four EEG microstate templates named MS1, MS2, MS3, and MS4 in the present work.

**Table 1.** The calculated evaluation indexes of the detected microstate templates after back-fitting across three experimental stages.

| Evaluation Index |  | MS1 | MS2 | MS3 | MS4 |
| --- | --- | --- | --- | --- | --- |
| Microstate Proportion (%) | pre-stimulus | 17.47 | 19.42 | 30.18 | 32.93 |
|  | video-stimulated | 18.91 | 18.93 | 30.62 | 31.55 |
|  | post-stimulus | 17.61 | 20.78 | 27.72 | 33.89 |
| Cronbach's $\alpha$ value | | 0.7312 | 0.7097 | 0.7131 | 0.7620 |

### 3.3 Results of Study 1

The activation differences of EEG microstates under different emotion states are examined in the perspectives of task effect, level effect, and evoking dynamics.

#### 3.3.1 Task effect

For emotion task effect analysis, we examine the activation difference of EEG microstates at pre-stimulus (before emotion-evoking task) and post-stimulus (after emotion-evoking task) stages. As shown in Fig. 7, a significant increasing trend (after FDR) is observed in MS2 coverage (*p* = 0.046) and MS4 coverage (*p* = 0.045), while a significant decreasing trend is observed in MS3 coverage (*p* = 0.005), duration (*p* = 0.029), and occurrence (*p* = 0.045). The transitions from MS3 to MS2 and from MS4 to MS2 significantly increase after emotion task manipulation, while the transition from MS4 to MS3 significantly decreases. These results reflect that the emotion-evoking task leads to a change in EEG microstate activities with distinct patterns. It is found that a positive task effect is observed in MS2 and MS4, meanwhile, a negative task effect is observed in MS3.

**Fig. 7.**
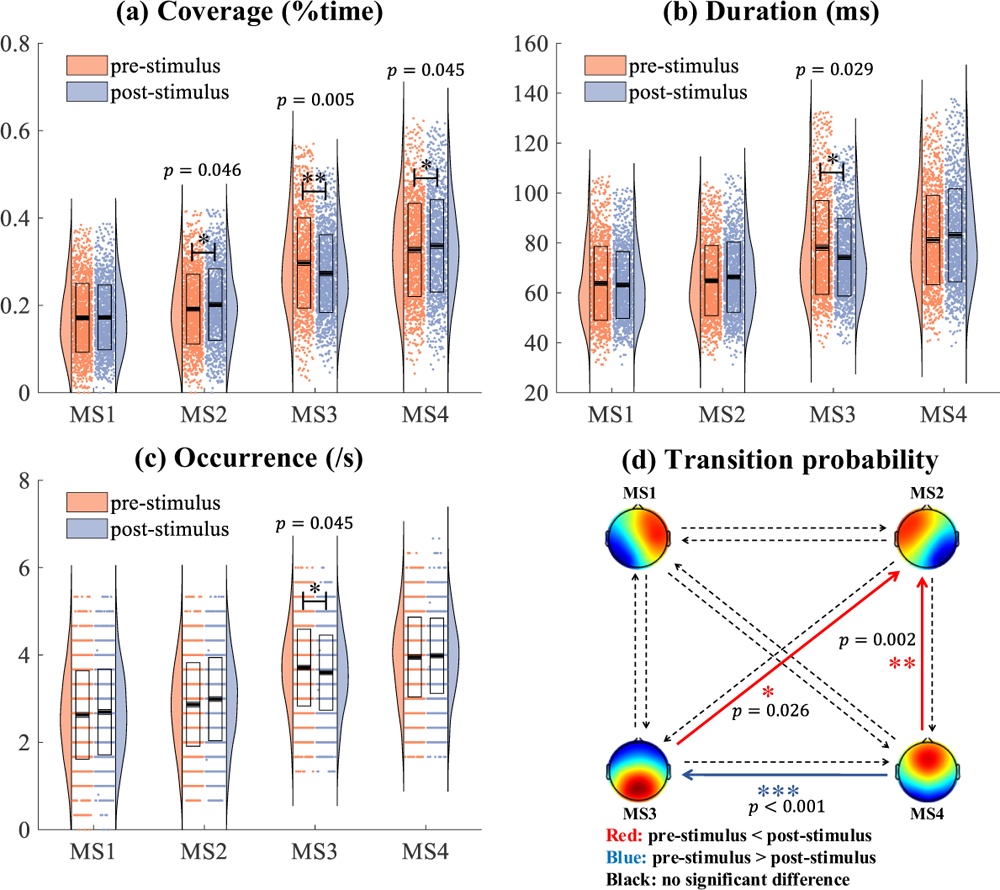
Emotion-evoking task effect evaluation on EEG microstate activities by measuring the paired difference between pre-stimulus and post-stimulus stages. The cross-subject statistical results of paired differences in terms of each microstate feature: (a) coverage, (b) duration, and (c) occurrence. Here, the dots represent the original feature distribution and the outlines of the violin plot represent the kernel probability density estimation. Box plots illustrate the inter-quartile ranges of the features, along with median lines in black. (d) The cross-subject statistical results of paired differences in terms of microstate transition. Red arrows indicate an increasing transition from pre-stimulus to post-stimulus, while blue arrows represent a decreasing transition after the emotion-evoking task occurs. All p-values are corrected with FDR, setting the statistical significance at 5%. (* p<0.05, ** p<0.01, *** p<0.001)

#### 3.3.2 Level effect

The influence of emotion levels (low/high) on brain responses is examined on two independent emotion dimensions (valence and arousal). For the valence dimension (Fig. 8), a lower MS4 occurrence is observed in the high valence group as compared to the low valence group, with the corresponding p-value of 0.038 (< 0.05). For the other EEG microstates (MS1, MS2, and MS3), no significant statistical difference is observed between low and high valence groups. By comparing the microstate transition probability between low and high valence groups, we observe a greater transition from MS1 to MS2 (*p* = 0.026) and a lower transition from MS1 to MS4 (*p* = 0.023) in the high valence group.

**Fig. 8.**
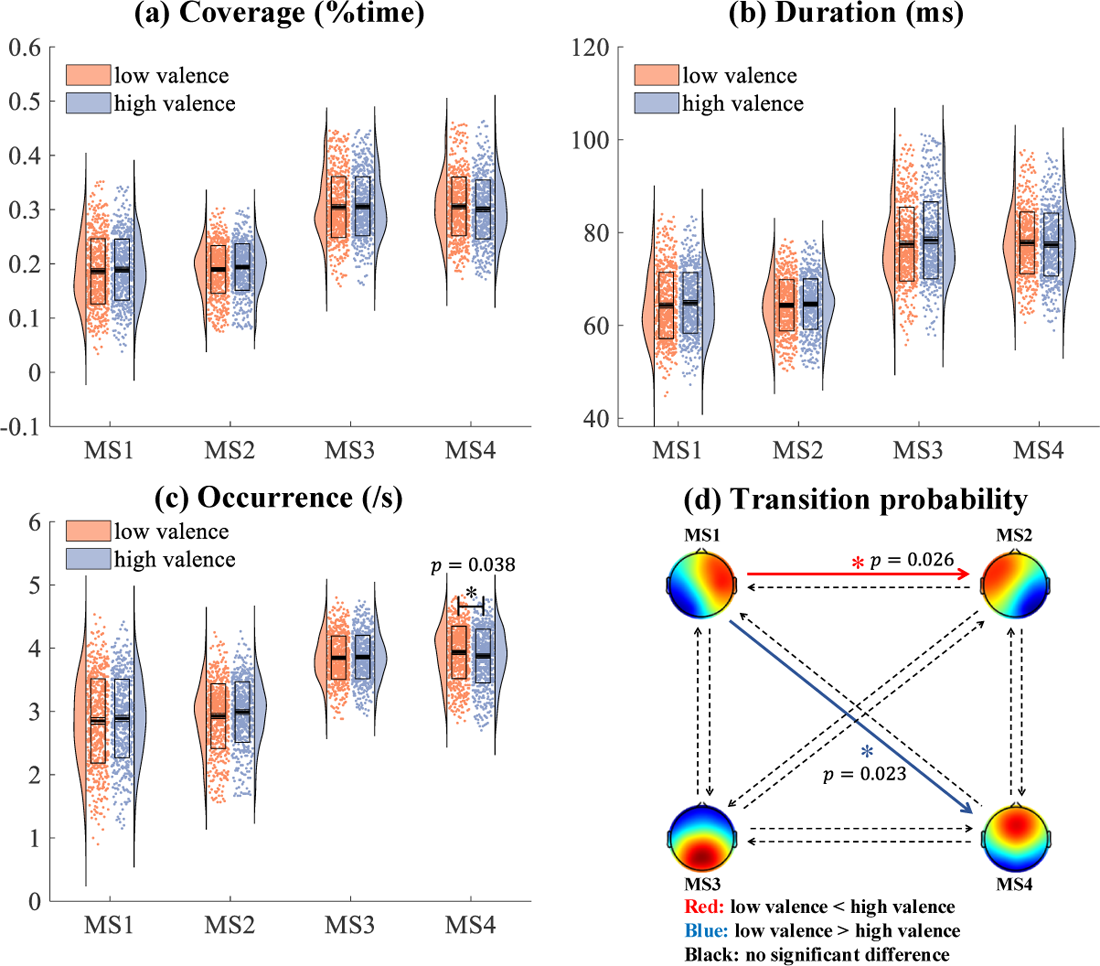
Valence-based level effect on EEG microstate activities. An independent t-test/Wilcoxon Rank Sum test is utilized to examine the statistical differences in EEG microstate activities between low and high valence groups. In the statistical results of (a) coverage, (b) duration, and (c) occurrence, the dots represent the original feature distribution and the outlines of the violin plot represent the kernel probability density estimation. Box plots illustrate the inter-quartile ranges of the features, along with median lines in black. In the statistical results of transition probability (d), red arrows indicate an increasing transition from low to high valence group, while blue arrows represent a decreasing transition in the high valence group as compared to the low valence group. (* p<0.05)

For the arousal dimension, the inter-group statistical differences between low and high arousal groups are shown in Fig. 9. It shows greater coverage (*p* = 0.0 5) and occurrence (*p* = 0.020) of MS3 are found in the high arousal group as compared to the low arousal group. For MS1, MS2, and MS4, no significant difference is observed between low and high arousal groups. By comparing the differences in microstate transition probability between low and high arousal groups, a higher transition probability from MS2 to MS3 (*p* = 0.024) is observed in the high arousal group.

**Fig. 9.**
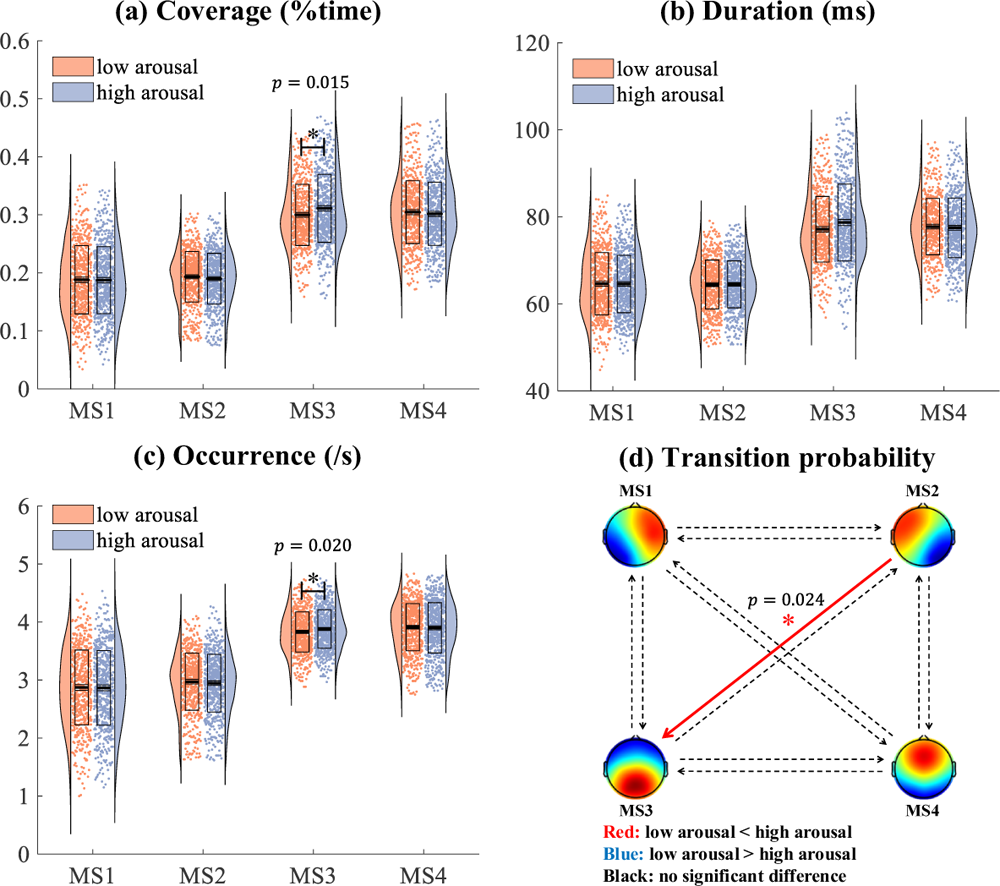
Arousal-based level effect on EEG microstate activities. An independent t-test/Wilcoxon Rank Sum test is utilized to examine the statistical differences in EEG microstate activities between low and high arousal groups. In the statistical results of (a) coverage, (b) duration, and (c) occurrence, the dots represent the original feature distribution and the outlines of the violin plot represent the kernel probability density estimation. Box plots illustrate the inter-quartile ranges of the features, along with median lines in black. In the statistical results of transition probability (d), red arrows indicate an increasing transition in the high arousal group compared to the low arousal group, while blue arrows represent a decreasing transition from the low to high arousal group. (* p<0.05)

The above results show that the level differences in emotion could be reflected by the patterns of microstate activities, especially MS3 and MS4. Distinct activation patterns of EEG microstates are observed on valence and arousal. MS4 is sensitive to the changes in valence levels, where high valence leads to a lower MS4 occurrence. In contrast, MS3 activity is related to arousal levels, where high arousal leads to higher MS3 coverage and occurrence.

#### 3.3.3 Evoking dynamics

The evoking emotion dynamics are measured as the EEG microstate activity differences between any two adjacent segments. These moment-to-moment changes are evaluated as the temporal variation of emotion-evoked EEG dynamics. In line with the previous observations that MS3 and MS4 play an important role in high-level cognitive function and conscious processing (Khanna et al. 2015), our results in the study of task effect and level effect also demonstrate that MS3 and MS4 are more related to emotion perception. Next, the exploration of emotion-related evoking dynamics will mainly focus on these two microstates (MS3 and MS4). For each trial, the evoking dynamics in terms of MS3 and MS4 features are analyzed across 32 subjects and a cross-subject multimedia-specific activation pattern is obtained for each video. Here, we take the coverage feature as an example and report the results in Fig. 10. As presented in Fig. 10 (a), varied activation patterns of EEG microstate responses in terms of MS3 and MS4 coverage are observed at 40 trials (videos). For each video, we carefully examine the temporal variations of EEG microstate activities. As shown in Fig. 10 (b), the time segments with significant statistical differences to the previous segments could be considered as “turning points” in the multimedia-evoked brain responses. According to our observations, different videos (different multimedia content) lead to different evoking patterns on the temporal characteristics of EEG microstate activities, where the turning points happen at different time moments and the temporal dynamic patterns of EEG microstate activities perform differently. A detailed observation about the evoking dynamics of video 20 is given as an example (Fig. 10 (c)). The turning points mainly occur from Seg 7 to Seg 12 (in total 20 segments; video length: 60 s; segment length: 3 s). For MS3 coverage, one turning point is observed from Seg 11 to Seg 12 with a significant decreasing trend. For MS4 coverage, significantly increasing trends are observed from Seg 7 to Seg 8, from Seg 9 to Seg 10, and from Seg 11 to Seg 12, and a significant decreasing trend is found from Seg 8 to Seg 9. The differences of the temporal dynamic evoking effects on emotion-related EEG microstate activities could be possibly explained by assuming that the content differences in multimedia stimulation would lead to different evoking reactions in the brain during emotion induction. More detailed evoking dynamic changes of each video are reported in Appendix IV of the Supplementary Materials.

**Fig. 10.**
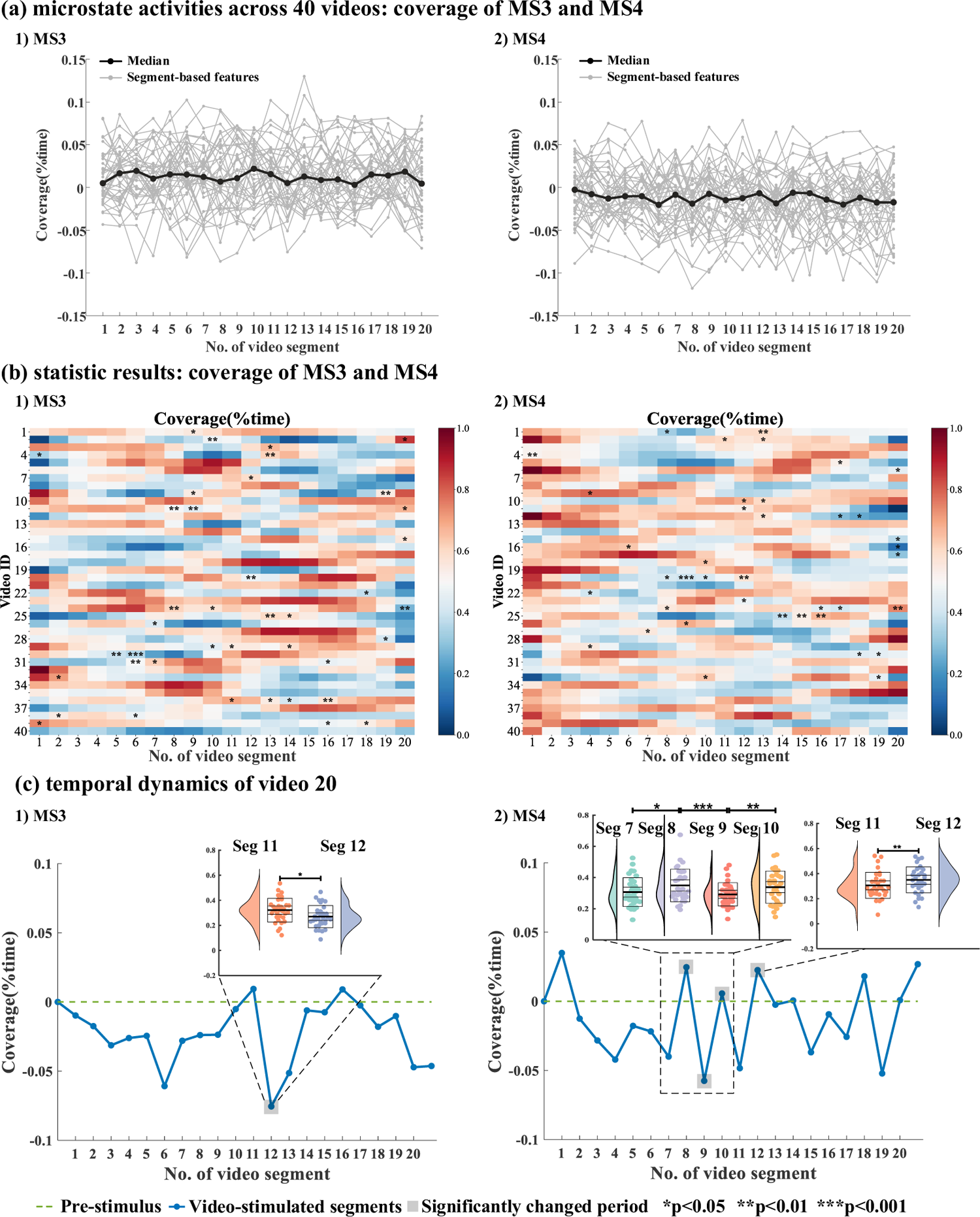
Video-evoked emotion dynamics in terms of segment-based EEG microstate activities. (a) Visualization of dynamic microstate activities in terms of MS3 and MS4 coverage across 40 videos. A gray line refers to one video’s temporal dynamic pattern and the black line is the median of all the temporal dynamic patterns across 40 videos. (b) Statistic results of the temporal variations of EEG microstate activities during emotion induction. In the heatmap, the colors refer to the microstate feature values, where the feature values are calculated as an average of segment-based MS3 or MS4 coverage features across 32 subjects for each video and normalized into the range of [0, 1] for visualization. The segments marked by “*”, “**”, or “***” refer to the turning points at which the microstate activities of the current time segment are significantly changed compared to the previous time segment. (c) A detailed example of the temporal changes in terms of MS3 and MS4 coverage under an emotion-evoking task using video 20. (* p<0.05, ** p<0.01, *** p<0.001)

Besides, the temporal distributions of the identified turning points for each video are reported in Table 2 (for MS3 coverage) and Table 3 (for MS4 coverage). It is found that the turning points are observed in 55% of videos (22/40). For MS3 coverage, it is found 22.7% of turning points are observed at the early stimulation stage, 54.5% at the middle stimulation stage, and 59.1% at the late stimulation stage. For MS4 coverage, 22.7% of turning points are found at the early stimulation stage, 50.0% at the middle stage, and 45.5% at the late stimulation stage. These results generally show that the turning points are mainly distributed at the middle and late stimulation stages, which suggests the temporal patterns of stimulation perception during emotion induction. Overall, the results obtained in Study 1 reveal that the dynamic changes of the evoked emotions can be well characterized by EEG microstate representations, especially by MS3 and MS4.

**Table 2.** The temporal distribution of turning points of 40 videos in terms of MS3 coverage.

| Video ID | Early stimulation<br>(Seg1-Seg8) | Middle stimulation<br>(Seg9-Seg14) | Late stimulation<br>(Seg15-Seg20) |
| --- | --- | --- | --- |
| 1 | x | √ | x |
| 2 | x | √ | √ |
| 3 | x | √ | x |
| 4 | x | √ | x |
| 5 | x | x | x |
| 6 | x | x | x |
| 7 | x | √ | x |
| 8 | x | x | x |
| 9 | x | √ | √ |
| 10 | x | x | x |
| 11 | x | √ | √ |
| 12 | x | x | x |
| 13 | x | x | x |
| 14 | x | x | x |
| 15 | x | x | √ |
| 16 | x | x | x |
| 17 | x | x | x |
| 18 | x | x | x |
| 19 | x | x | x |
| 20 | x | √ | x |
| 21 | x | x | x |
| 22 | x | x | √ |
| 23 | x | x | x |
| 24 | x | √ | √ |
| 25 | x | √ | √ |
| 26 | √ | x | x |
| 27 | x | x | x |
| 28 | x | x | √ |
| 29 | x | √ | √ |
| 30 | √ | x | x |
| 31 | √ | x | √ |
| 32 | x | x | x |
| 33 | √ | x | x |
| 34 | x | x | x |
| 35 | x | x | x |
| 36 | x | √ | √ |
| 37 | x | x | x |
| 38 | √ | x | x |
| 39 | x | x | √ |
| 40 | x | x | √ |

**Table 3.** The temporal distribution of turning points of 40 videos in terms of MS4 coverage.

| Video ID | Early stimulation<br>(Seg1-Seg8) | Middle stimulation<br>(Seg9-Seg14) | Late stimulation<br>(Seg15-Seg20) |
| --- | --- | --- | --- |
| 1 | x | √ | x |
| 2 | x | √ | x |
| 3 | x | x | x |
| 4 | x | x | x |
| 5 | x | x | √ |
| 6 | x | x | √ |
| 7 | x | x | x |
| 8 | x | x | x |
| 9 | √ | x | x |
| 10 | x | √ | x |
| 11 | x | √ | x |
| 12 | x | √ | √ |
| 13 | x | x | x |
| 14 | x | x | x |
| 15 | x | x | √ |
| 16 | √ | x | √ |
| 17 | x | x | √ |
| 18 | x | √ | x |
| 19 | x | x | x |
| 20 | x | √ | x |
| 21 | x | x | x |
| 22 | √ | x | x |
| 23 | x | √ | x |
| 24 | x | √ | √ |
| 25 | x | x | √ |
| 26 | x | √ | x |
| 27 | √ | x | x |
| 28 | x | x | x |
| 29 | √ | x | x |
| 30 | x | x | √ |
| 31 | x | x | x |
| 32 | x | x | x |
| 33 | x | √ | √ |
| 34 | x | x | x |
| 35 | x | x | x |
| 36 | x | x | x |
| 37 | x | x | x |
| 38 | x | x | x |
| 39 | x | x | x |
| 40 | x | x | x |

### 3.4 Results of Study 2

To characterize the stimulation effect of multimedia content on the patterns of microstates, we separately analyze the temporal correlation between EEG microstate activities and multimedia stimulation in terms of visual content, audio content, and timing effect.

#### 3.4.1 Visual content

The correlations between EEG microstate activities and visual content in terms of low-level and high-level visual features are reported in Fig. 11 (a) and (b). The results show that the changes in visual content have a close relationship with the dynamic activities of EEG microstates, which is mainly reflected in the MS4 activity. For low-level visual features, a positive correlation is found between the brightness and MS4 occurrence, where a higher MS4 occurrence is observed when the presented videos with high brightness. For high-level visual features, a positive correlation is observed between the visual features extracted from *conv*11 to *conv*13 and MS4 coverage and duration. Besides, a negative association is also observed between the high-level visual features extracted from *conv*11 to *conv*12 and MS3 coverage. Compared to the stimulation effect of low-level visual features, a more complex stimulation effect is found for high-level visual features, suggesting that high-level features are more related to the changing activities of MS3 and MS4 and play a more important role in emotion induction.

**Fig. 11.**
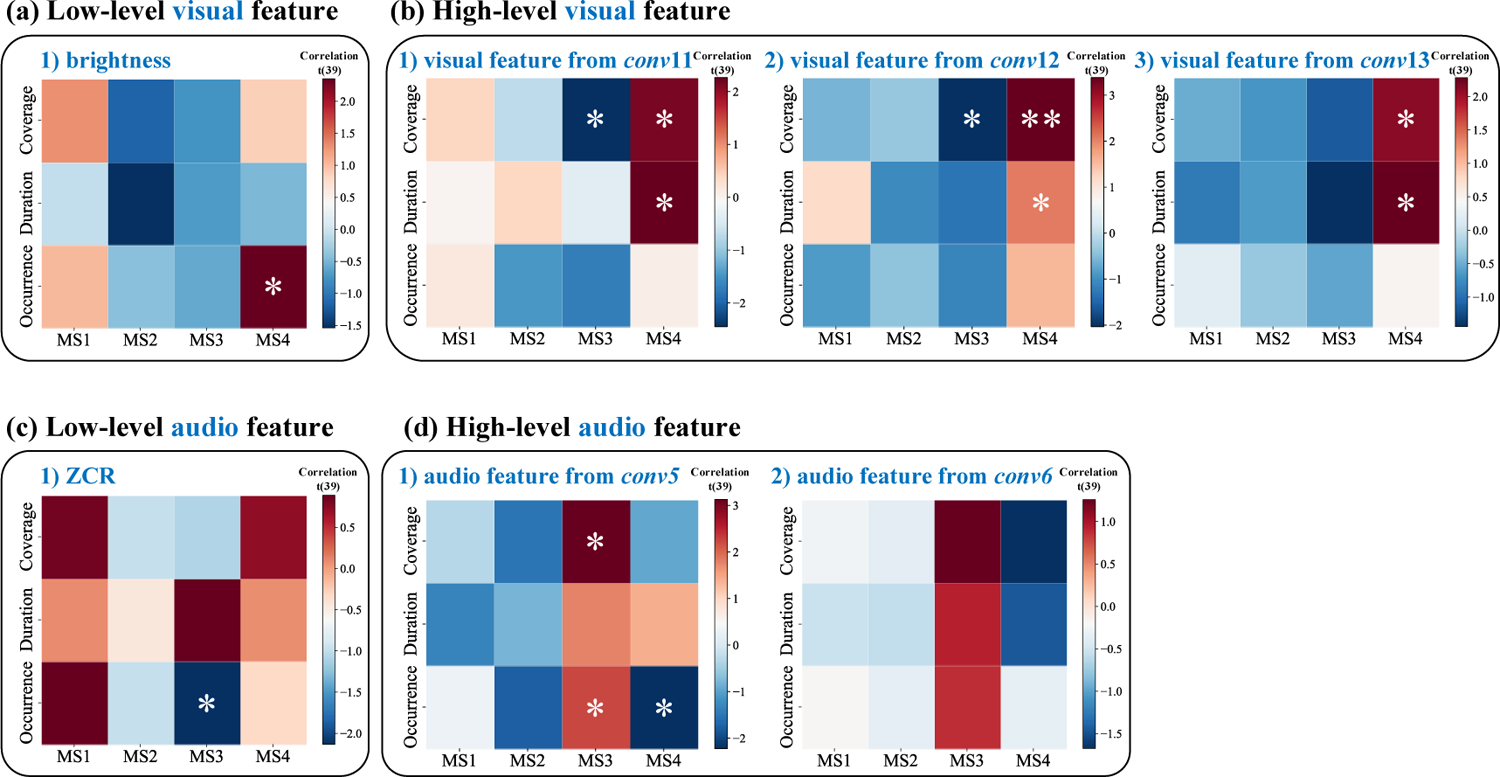
Stimulation effect of visual and audio content on segment-based microstate features. The heatmap is a visualization display of the calculated t values from correlation relationship measurement. A positive t value is marked as red, which indicates a positive correlation between stimulation content and EEG microstate activities. A negative t value is marked as blue, which refers to a negative correlation between stimulation content and microstate activities. (* p<0.05, ** p<0.01)

#### 3.4.2 Audio content

Similar to the visual content effect analysis in Section 3.4.1, the correlations between audio content and EEG microstate activities are reported in Fig. 11 (c) and (d). The results show that audio content mainly influences MS3 activity. For low-level audio features, a negative correlation (an increase of ZCR feature leads to a decrease of MS3 occurrence) is observed. For high-level audio features, a positive correlation (the audio features extracted from *conv*5 activate MS3 responses for a larger coverage and occurrence) is found. At the same time, a negative correlation is observed between the features extracted from *conv*5 and MS4 occurrence. In the investigation of the audio content effect in terms of high-level audio features, a complex correlation between EEG microstate activities of MS3 and MS4 and the high-level audio features extracted from *conv5* is observed. However, for the high-level audio features extracted from *conv*6, no significant correlation with EEG microstate activities is found. One possible reason could be that the utilized VGGish was trained for audio classification tasks based on the Youtube-8M database (Abu-El-Haija et al. 2016), and the audio features characterized at the last convolutional layer (*conv*6) could reflect more about classification information instead of the affective information.

#### 3.4.3 Timing effect

We shift the multimedia stimulation content in a preceded or succeeded direction with a time range of −1s to 1s and measure the corresponding temporal correlation with the EEG microstate activities at every time-shifting parameter. For visual content perception, the results (Fig. 12 (a)) reveal that the valid time effect is in the range of −100 ms to 400 ms. The highest correlation is reached at 0 ms (stimulation onset), and then the correlation declines from 0 ms to 400 ms. Besides, a preceded effect of visual content on EEG microstate dynamics is also observed before the stimulation onset (−100 ms). One possible reason could be that there may exist an expectation effect before visual content presentation, as the adjacent stimulation content in continuous videos is closely content related. For audio content perception, it is found that the timing effect mainly occurs from 0 to 600 ms (Fig. 12 (b)). The highest correlation happens at 200 ms, which shows a post-stimulus effect of audio content on brain responses during emotion induction. These results show that, for visual and audio content, the timing effects on simultaneous brain responses are different, which results in a distinct activation pattern of EEG microstate activities during multimedia-evoked emotion induction. Stimulation perception of visual content is closely related to MS4 activities with an onset effect, and that of audio content is more related to MS3 activities with a post-stimulus effect.

**Fig. 12.**
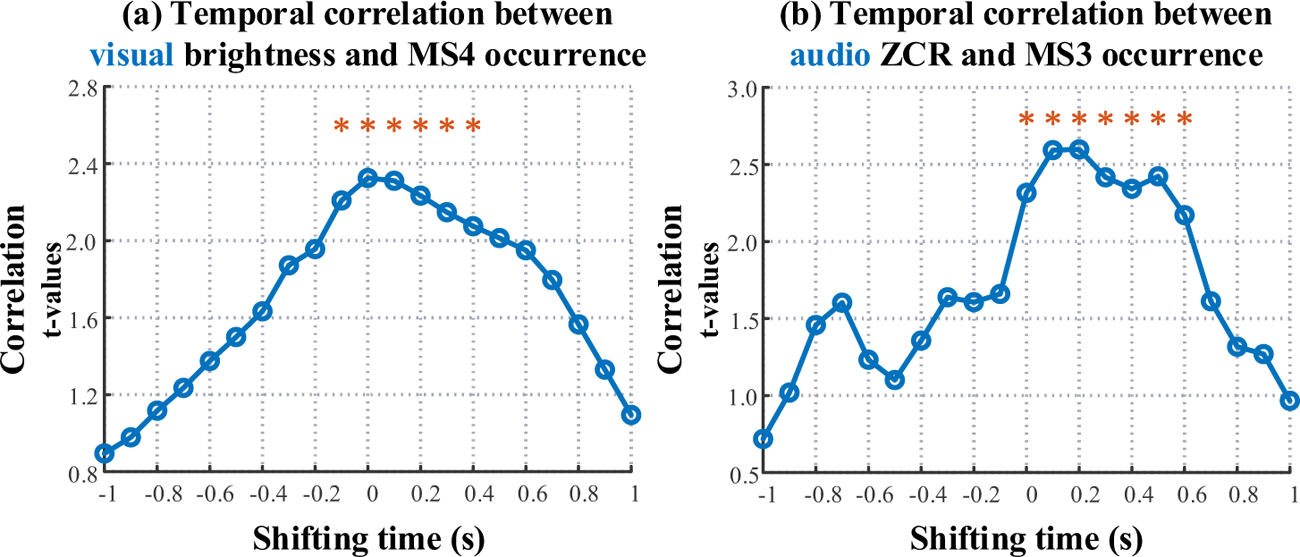
Temporal correlation between shifted multimedia content and EEG microstate activities. (a) Temporal correlation between the time-shifting visual content (brightness) and MS4 occurrence. (b) Temporal correlation between the time-shifting audio content (ZCR) and MS3 occurrence. (* p<0.05; ZCR: zero-crossing rate)

## 4 Discussion

The main goal of this study is to discover the associations among the appearance of microstates, emotion dynamics, and stimulation content. Both the evoked emotions and the used stimulation for emotion-evoking are analyzed and the corresponding specific parameters of EEG microstates that are significantly changed are identified. The results show, functioning as an informative intermediary, EEG microstates map the subjective emotion states and present multimedia stimulation content to dynamic activity patterns of the brain. The observations of EEG microstate dynamics in video-evoked emotion study and the potential neural mechanisms underlying emotion perception are further discussed in this section.

### 4.1 Neurophysiological significance of EEG microstates

In this work, four EEG microstate templates are detected using a sequential clustering method. It is found that the identified EEG microstates share similar topographical configurations to the canonical EEG microstates in the previous resting-state EEG studies (Britz et al. 2010; Musso et al. 2010; Khanna et al. 2015; Michel and Koenig 2018). According to the literature, MS1, MS2, MS3, and MS4 are functionally mapped to four important functional brain networks (Table 4), including the auditory network, visual network, default mode network (DMN), and dorsal attention network (DAN). For example, Britz et al. (2010) validated the spatial correlation between EEG microstates and the four functional brain networks based on the simultaneous EEG-fMRI data. In their study, it was detected that MS1 was spatially correlated with the negative BOLD activation over bilateral superior and middle temporal gyri, MS2 was mapped with the negative BOLD activation in bilateral extrastriate visual areas, MS3 was related with the positive BOLD signals in the anterior cingulate cortex and bilateral inferior frontal gyri that are important brain regions for emotion-related salience information integration, and MS4 was correlated with the negative BOLD activation in the right-lateralized dorsal and ventral areas of frontal and parietal cortex (these brain regions functionally activate for attention shifting and reorienting task during external stimulation processing). Besides, through the source localization of identified EEG microstates across 164 subjects, Custo et al. (2017) found the corresponding activation relationships between cortical regions and EEG microstates were: the source localization of MS1 laid in the left middle and superior temporal lobe and the left insular cortex; the source localization of MS2 mainly laid in the left and right occipital cortices; the source localization of MS3 laid in the precuneus and the posterior cingulate cortex; and the source localization of MS4 laid in the right inferior parietal lobe, the right middle, and superior frontal gyri. These spatial analyses of the neuronal generators of EEG microstates through fMRI analysis or source localization supports the neurophysiological functional association between EEG microstates and specific large-scale brain networks.

**Table 4.**
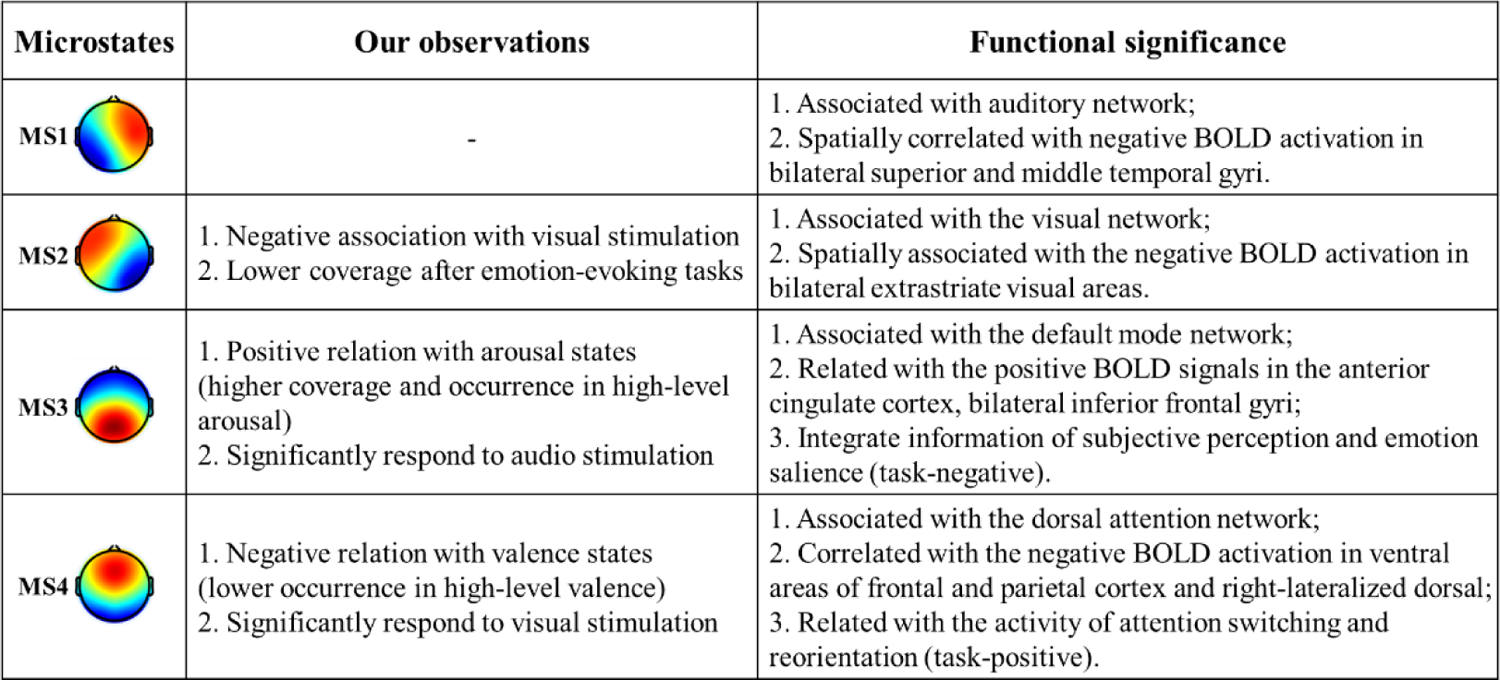
A summary of the neurophysiological significances of EEG microstates.

| Microstates | Our observations | Functional significance |
| --- | --- | --- |
| MS1 | - | 1. Associated with auditory network;<br>2. Spatially correlated with negative BOLD activation in bilateral superior and middle temporal gyri. |
| MS2 | 1. Negative association with visual stimulation<br>2. Lower coverage after emotion-evoking tasks | 1. Associated with the visual network;<br>2. Spatially associated with the negative BOLD activation in bilateral extrastriate visual areas. |
| MS3 | 1. Positive relation with arousal states (higher coverage and occurrence in high-level arousal)<br>2. Significantly respond to audio stimulation | 1. Associated with the default mode network;<br>2. Related with the positive BOLD signals in the anterior cingulate cortex, bilateral inferior frontal gyri;<br>3. Integrate information of subjective perception and emotion salience (task-negative). |
| MS4 | 1. Negative relation with valence states (lower occurrence in high-level valence)<br>2. Significantly respond to visual stimulation | 1. Associated with the dorsal attention network;<br>2. Correlated with the negative BOLD activation in ventral areas of frontal and parietal cortex and right-lateralized dorsal;<br>3. Related with the activity of attention switching and reorientation (task-positive). |

Given the observed neurophysiological significances of EEG microstates, researchers also applied EEG microstate analysis for brain dynamics decoding. For example, Gui et al. (2020) estimated the language processing of unresponsive patients by mapping MS1 and MS2 into an Anterior-Posterior (A-P) map and mapping MS3 and MS4 into a Left-Right (L-R) map. They observed a higher activation of an A-P state (MS1 and MS2) but a lower activation of the L-R state (MS3 and MS4) in patients when compared to the healthy group. The results support that MS1 and MS2 mainly respond to low-level sensory brain processing, whereas MS3 and MS4 respond to high-level cognitive information manipulation. The activation bias between low-level sensory processing and high-level cognitive information manipulation would be used to explain the results of the patients with the disorders of consciousness. Hence, taking the functional significance of EEG microstates in neurophysiology into consideration, the underlying neural mechanism of emotions could be further revealed by inspecting the functional patterns of EEG microstate activities.

### 4.2 Microstates as indicators of emotion states

The neural processing mechanisms of emotion perception in the brain can be inferred from the relationship between EEG microstate activities and subjective-experienced emotion states. First, in task effect, the statistical differences in EEG microstate activities between pre-stimulus and post-stimulus stages show that emotional task manipulations lead to an increase in MS2 and MS4 coverage and a decrease in MS3 coverage, duration, and occurrence. The representation difference in task effect can be possibly explained that the fast-changing visual stimulation in an emotional video activates the visual network increasing MS2 activities (Milz et al. 2016). For the decrease of MS3 activities, one possible reason could be video-watching emotion inhibit the activation of the DMN in line with the findings that the DMN is task-negative that mainly relates to intrinsic information processing like memory recall, self-judgments, prospective thinking (Britz et al. 2010; Yuan et al. 2012; Seitzman et al. 2017). The review of Satpute et al. (2019) highlighted the functional role of DMN in emotion perception by conceptualizing the stimulation content (Gu et al. 2013). The increase of MS4 activities is in line with the functional activity of the DAN (task-positive) that would be activated in multimedia-directed emotion induction for attention reorientation and focus switching (Nummenmaa et al. 2012; Szczepanski et al. 2013). However, no significant difference is found in MS1 (related to the auditory network). According to Britz et al.’s work (2010), it was found that MS1 simultaneously corresponded to a negative BOLD activation mainly in bilateral superior and middle gyri (auditory-related functional brain regions) and the primary visual cortex of the bilateral extrastriate cortex (visual-related functional brain regions) (Mantini et al. 2007). In Milz et al.’s work for cognitive task-based EEG microstate analysis, a larger MS1 duration was found during the visual-stimulated task compared to the resting-state and audio-stimulated task (Milz et al. 2016). In our case, as video-based multimedia stimulation is adopted for emotion induction, the simultaneous presentation of audiovisual content would possibly inhibit the activities of MS1.

Additionally, the significant differences in MS3 and MS4 activities during emotion induction may indicate that MS3 and MS4 accompanied with the DMN and DAN are more related to the high-level perceptual processing of emotion in the brain (Morawetz et al. 2016; Iordan and Dolcos 2017; Satpute and Lindquist 2019). Similar to the previous works, for example, Seitzman et al. (2017) found that task manipulation mainly altered the activities of MS3 and MS4 with an inhibiting effect on MS3 but a promoted effect on MS4. For serial subtraction tasks, the coverage, duration, and occurrence of MS3 significantly decreased and these features of MS4 significantly increased after task manipulation. Similar results were also observed by Kim et al. (2021) that a significant decrease of MS3 and a significant increase of MS4 were found in good performance groups while performing mental arithmetic tasks. These findings generally suggest that the influences of cognitive task manipulation (refers to emotion induction in this work) on microstate activities are in line with task effects on the DMN (task-negative, associated with MS3) and DAN (task-positive, associated with MS4).

In the investigation of microstate representation differences on valence and arousal dimensions, we also observe that emotion level differences mainly influence the activities of MS3 and MS4. Valence level is positively correlated with MS4 occurrence, while arousal level is negatively correlated with MS3 coverage and occurrence. These findings are congruent with the observations of emotion level effects on the activities of functional brain networks in the previous studies. Nummenmaa et al. (2012) found that low valence increased the activities of the DAN for emotion perception and induction, while high arousal activated the DAN for attention switching via external multimedia stimulation. Besides, Colibazzi et al. (2010) obtained similar observations in a self-generated emotion induction task. For low valence emotions, higher BOLD signals were detected in the right dorsolateral prefrontal cortex and rostral dorsal anterior cingulate cortex that are spatially belonged to the DAN. For high arousal emotions, higher BOLD responses were detected in the midline and medial temporal lobes that are belonged to the DMN. All the above results suggest that the DAN and DMN are essential for emotion processing, where DAN (MS4) mainly functions for valence-related emotion processing, and the DMN (MS3) functions for arousal-related emotion processing.

Furthermore, the representation differences of EEG microstate activities on valence and arousal dimensions can be interpreted by the functional significances of the DAN and DMN during emotion induction. As well investigated in the literature, the DAN plays a dominant role in stimulus-directed sensory and functions for extracting emotion-related salient information for valence induction (Szczepanski et al. 2013). The DMN is mainly involved in self-reflection and internal perception, both of which are important for arousal processing (McKiernan et al. 2003). These findings are also reflected in our observations of the emotion level effect on EEG microstate activities, resulting in a positive association between valence level and MS4 activities and a negative association between arousal level and MS3 activities.

### 4.3 Microstates and stimulation content

In this work, the temporal association between multimedia content and EEG microstate activities is measured to describe the dynamic process of emotion perception in the brain. In previous affective computing studies, it was found that visual brightness as a fundamental and commonly used visual feature has been validated as an essential visual attribute for emotional valence induction in video affective content analysis (Wang and Ji 2015). For the audio ZCR feature, it has been validated as a key acoustic feature for emotion arousal enhancement and has been widely applied for multimedia content-based emotion recognition (Zhang et al. 2010). In our study, these multimedia features are also found to be highly associated with emotion-evoked EEG microstate activities that a higher brightness leads to a higher occurrence of MS4, and a higher ZCR leads to a lower MS3 occurrence. The stimulation effect is mainly found in MS3/ MS4 activities with low-level audio/visual features instead of MS1/MS2 activities. One possible reason would be that emotion induction is a complex perceptual process associated with a subjective understanding of presented stimulation content, where the low-level visual and audio features could also be considered as important cues of subjective-experienced emotions. These observations are consistent with the findings presented in (Kurt et al. 2017; Seng et al. 2018).

Through investigating the temporal correlation between multimedia stimulation and dynamic EEG activities during emotion induction, our findings demonstrate that emotion perception is a temporally dynamic process coordinated with the stimulation of multimedia content (Effron et al. 2006). For visual content, the temporal association between visual content and dynamic EEG activities is observed in the time range of −100–400 ms. These results are consistent with the previous findings. For example, Oya et al. (2002) observed a strong gamma response around 150– 450 ms on the amygdala under the visual task using emotional pictures. In Potts et al.’s visual-stimulated ERP task (Potts 2004), it was found the P2a (about 220–316 ms) component significantly increased in response to the visual content. In this work, a preceded correlation (–100 ms) is found between visual content and the triggered emotions, which could be explained by the fact that emotion induction is a temporally dynamic process and the previous stimulation continuously affects the following emotion states. Through exploring the potential associations between continuous multimedia content and dynamic EEG microstate activities, it helps us to understand the temporal characteristics of video-evoked emotion processing in the brain.

To further understand the emotion modulated neural responses, we examine the temporal association between EEG microstate activity dynamics and continuous stimulation of multimedia content in terms of the timing effect. Considering the EEG microstate representations in terms of evoking dynamics (Section 3.3.3) and timing effect (Section 3.4.3), we can infer that emotion induction is a temporally dynamic process reflected in the temporal coordination between EEG microstate activities and the objective-presented stimulation content during video-watching emotion induction tasks.

### 4.4 Limitations and future work

There remains a lack of clarified identification of emotion-specific EEG microstates. The specialized roles of EEG microstate activities in interpreting the dynamic process of emotion processing and regulation are still an open issue. The extension of applying EEG microstate analysis into video-triggered emotion induction study offers a novel perspective into emotion dynamics, as well as border knowledge of neurophysiological significances of EEG microstates. Besides, there is still a lack of locating surface EEG signals of emotion-related EEG microstates into the corresponding neuronal activity. It will limit the interpretability of EEG microstate representation patterns in dynamic emotion analysis. To extend, EEG microstate analysis with voxel-based source localization would greatly enhance the spatial resolution for accurate reflection of global neuronal activities and improve the performance of functional brain activity estimation, monitoring, and regulation. Moreover, fMRI data with high spatial resolution can capture the hemodynamic changes in deep brain structures. Combining EEG microstates and fMRI data will provide a more comprehensive investigation of emotion-related neurophysiological mechanisms in high temporal and spatial resolution.

Additionally, instead of conducting video segmentation with fixed segment length, an adaptive content-based segmentation method would offer more reliable clues into the investigation of emotion perceptual mechanisms during video watching. In the current work, we simply segment the time process of video watching into the early, middle, and late stimulation stages with a fixed time length. The EEG microstate activity differences among different stimulation stages are measured for evoking dynamics analysis. However, different stimulation content would yield a varied temporal pattern of emotion-evoking. Taking advantage of the development in the computer vision field, content-based video segmentation methods like explicit and implicit video affective content analysis (Zhang et al. 2010; Wang and Ji 2015) could be incorporated for a better inspection into the neural mechanisms of emotional dynamics.

## 5 Conclusion

In summary, our work mainly focuses on the exploration of EEG microstates patterns in associations with emotion dynamics. A mapping among subjective-experienced emotion states, objective-presented stimulation content, and microstates is measured, and the use of EEG microstates to reveal potential emotion relevance is explored. The results show EEG microstates are capable of representing the dynamic characteristics of emotion-related EEG dynamics in continuous video-triggered emotion induction tasks. Especially, MS3 and MS4, which have a close association to high-level functional brain networks, show a good performance in the studies of subjective-experienced emotion state analysis and objective-presented stimulation effect analysis. Distinctive EEG microstate activity patterns are observed under different emotion states from the perspectives of task effect, level effect, and timing effect. Also, it is found that the emotional stimulation effect on EEG microstate patterns can be inferred by the objective-presented video content. This work of applying EEG microstate analysis into dynamic emotion analysis borders our knowledge of the functional significance of EEG microstates and provides an attractive approach for time-varied emotion-related neural mechanism exploration.

## Acknowledgments

This work was supported in part by the National Natural Science Foundation of China under Grant 61906122, in part by Shenzhen-Hong Kong Institute of Brain Science-Shenzhen Fundamental Research Institutions (2021SHIBS0003), in part by the Tencent “Rhinoceros Birds”-Scientific Research Foundation for Young Teachers of Shenzhen University, and in part by the High Level University Construction under Grant 000002110133.

## Supplementary Materials

## Appendix I: DEAP Database and EEG Processing

In this work, the public DEAP database with 32 subjects’ EEG recordings is used to explore the emotion modulation effect on EEG microstate representation patterns and transitions. The DEAP database is a well-known EEG-emotion dataset for human affective study (Koelstra et al. 2012). Forty music videos with a fixed length of 60 seconds were presented in a random sequence to every subject for emotion-evoking. Simultaneously, 32-electrode EEG signals were recorded at a sampling rate of 512 Hz using a Biosemi ActiveTwo system, where the electrodes were placed based on the international 10-20 system. Every single trial included four parts: pre-stimulus, video-stimulated, post-stimulus, and self-assessment. At the pre-stimulus and post-stimulus stages, subjects were instructed to look at the fixation cross located at the center of the screen with a relaxed and clear mind. At the video-stimulated stage, subjects were instructed to focus on watching the presented video for emotion induction. Subsequently, a self-assessment manikin (SAM) (Morris 1995) with a continuous scale from 1 to 9 was utilized to collect the self-assessment feedback on evoked emotions from the dimensions of valence (related with pleasantness) and arousal (related with excitation). The experimental pipeline of the DEAP database is shown in Fig. S1.

**Fig. S1.**
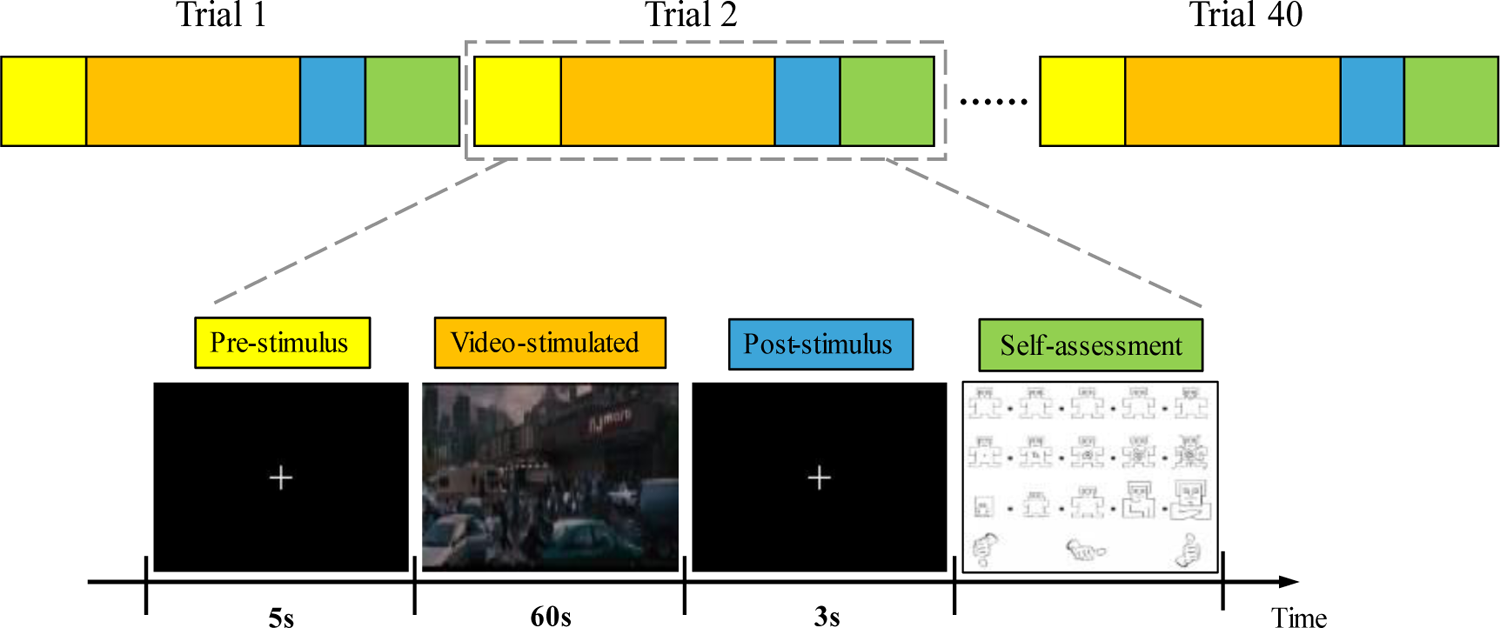
The experimental pipeline of the DEAP database.

Based on the collected raw EEG signals, a standard preprocessing pipeline is conducted for unwanted noise removal. Firstly, each trial of EEG recordings is filtered with a bandpass filter of 1 - 45 Hz for unrelated artifact filtering and a notch filter of 50 Hz for power line noise removal. Secondly, noisy electrodes are interpolated using an average strategy with the neighboring 3 electrodes. Thirdly, the filtered EEG data are adjusted to spatial zero-mean distribution by conducting a common average re-reference (Brunet et al. 2011). The common average referencing is implemented not only to remove zero-mean random noise for data quality improvement but also to adjust the preprocessed EEG data to a reference-free electric potential distribution for further spatial similarity detection (Pascual-Marqui et al. 1995; Murray et al. 2008). Finally, independent component analysis (ICA) is implemented for ocular artifact removals, such as eye movements and blinks. Meanwhile, other artifacts caused by body movement, cardiac or muscular activity are rejected as much as possible by manual screening. After the preprocessing, the noises in the EEG raw data are greatly removed and the clean EEG data are reconstructed by using the remained independent components. Notably, to minimize the impact of evoked emotions in the previous trial, we only used the last 3 s of the original 5 s pre-stimulus data for further analysis in this paper. Then, EEG data segmentation is conducted to separate each single-trial data into three stages: **pre-stimulus (3 s), video-stimulated (60 s), and post-stimulus(3 s)**.

## Appendix II: Subject-Representative Microstate Topographies

A detailed illustration of the obtained subject-representative microstate topographies from the three experimental stages at the first-step clustering of sequential microstate detection is given in Fig. S2. For the subject-representative microstate detection of each subject, the candidate topographies of the 40 trials of EEG data under pre-stimulus, video-stimulated, or post-stimulus stages are separately extracted from the local maximal of the GFP peaks, using a modified k-means clustering algorithm (Pascual-Marqui et al. 1995). The corresponding class numbers and GEV values of subject-representative microstates are reported in Table S1.

**Fig. S2.**
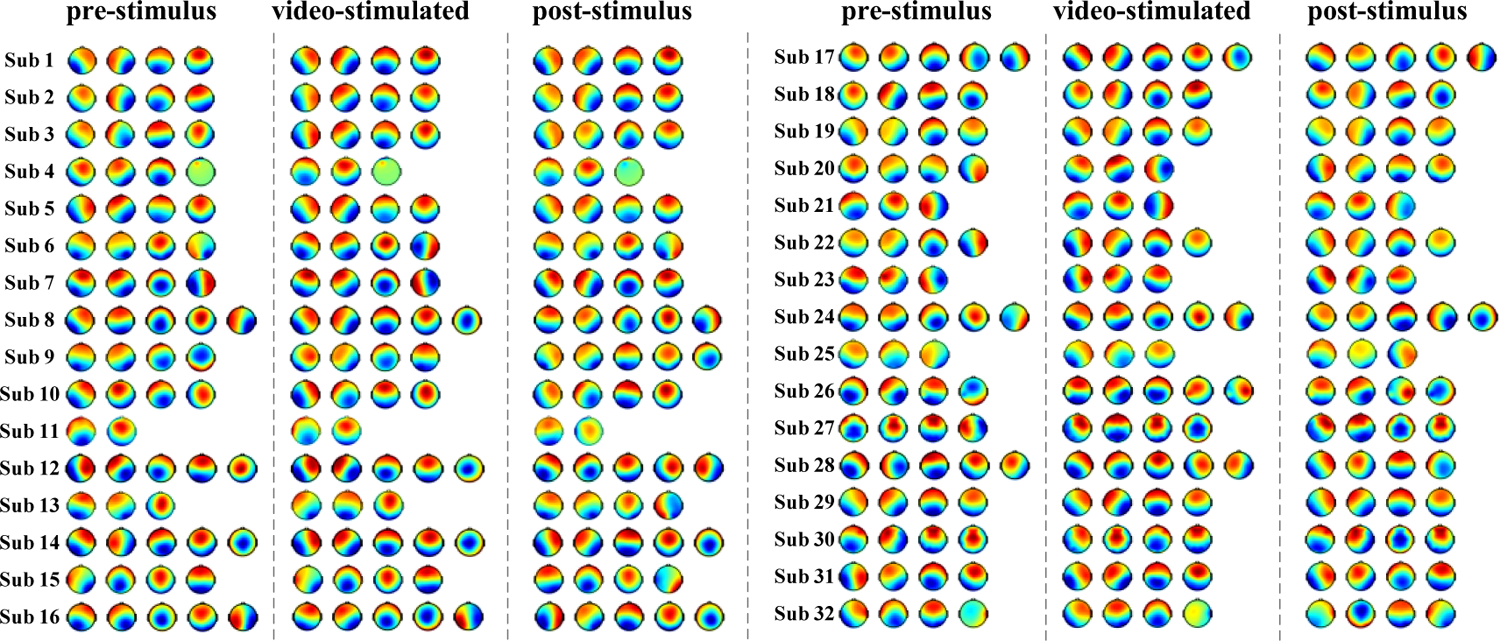
The topographical illustration of the obtained subject-representative microstate topographies extracted from the three experimental stages (pre-stimulus, video-stimulated, and post-stimulus) at the first-step clustering of the sequential microstate detection (within-subject-level clustering).

**Table S1.** The class numbers and GEV values of the obtained subject-representative microstates at the first clustering step of the sequential microstate detection.

| Subject ID | pre-stimulus |  | video-stimulated |  | post-stimulus |  |
| --- | --- | --- | --- | --- | --- | --- |
| | Class<br>Number | GEV<br>(mean ± std) | Class<br>Number | GEV<br>(mean ± std) | Class<br>Number | GEV<br>(mean ± std) |
| S1 | 4 | 0.72 ± 0.03 | 4 | 0.71 ± 0.02 | 4 | 0.72 ± 0.03 |
| S2 | 4 | 0.68 ± 0.04 | 4 | 0.69 ± 0.01 | 4 | 0.68 ± 0.04 |
| S3 | 4 | 0.68 ± 0.03 | 4 | 0.70 ± 0.01 | 4 | 0.70 ± 0.03 |
| S4 | 4 | 0.62 ± 0.08 | 3 | 0.60 ± 0.07 | 3 | 0.59 ± 0.09 |
| S5 | 4 | 0.74 ± 0.04 | 4 | 0.72 ± 0.03 | 4 | 0.76 ± 0.05 |
| S6 | 4 | 0.72 ± 0.03 | 4 | 0.70 ± 0.02 | 4 | 0.72 ± 0.04 |
| S7 | 4 | 0.66 ± 0.04 | 4 | 0.64 ± 0.03 | 4 | 0.65 ± 0.04 |
| S8 | 5 | 0.71 ± 0.03 | 5 | 0.71 ± 0.02 | 5 | 0.70 ± 0.04 |
| S9 | 4 | 0.74 ± 0.04 | 4 | 0.69 ± 0.02 | 5 | 0.75 ± 0.04 |
| S10 | 4 | 0.66 ± 0.05 | 4 | 0.65 ± 0.03 | 4 | 0.64 ± 0.04 |
| S11 | 2 | 0.51 ± 0.06 | 2 | 0.52 ± 0.06 | 2 | 0.52 ± 0.07 |
| S12 | 5 | 0.68 ± 0.03 | 5 | 0.68 ± 0.02 | 5 | 0.68 ± 0.03 |
| S13 | 3 | 0.65 ± 0.03 | 3 | 0.68 ± 0.03 | 4 | 0.67 ± 0.03 |
| S14 | 5 | 0.71 ± 0.03 | 5 | 0.71 ± 0.01 | 5 | 0.71 ± 0.03 |
| S15 | 4 | 0.66 ± 0.05 | 4 | 0.67 ± 0.03 | 4 | 0.66 ± 0.04 |
| S16 | 5 | 0.66 ± 0.04 | 5 | 0.65 ± 0.03 | 5 | 0.65 ± 0.04 |
| S17 | 5 | 0.70 ± 0.03 | 5 | 0.71 ± 0.02 | 5 | 0.73 ± 0.04 |
| S18 | 4 | 0.68 ± 0.05 | 4 | 0.67 ± 0.03 | 4 | 0.69 ± 0.07 |
| S19 | 4 | 0.69 ± 0.03 | 4 | 0.70 ± 0.01 | 4 | 0.70 ± 0.03 |
| S20 | 4 | 0.75 ± 0.04 | 3 | 0.66 ± 0.02 | 4 | 0.73 ± 0.05 |
| S21 | 3 | 0.63 ± 0.03 | 3 | 0.63 ± 0.02 | 3 | 0.63 ± 0.03 |
| S22 | 4 | 0.69 ± 0.05 | 4 | 0.69 ± 0.03 | 4 | 0.68 ± 0.05 |
| S23 | 3 | 0.61 ± 0.06 | 3 | 0.65 ± 0.04 | 3 | 0.61 ± 0.06 |
| S24 | 5 | 0.74 ± 0.03 | 5 | 0.72 ± 0.01 | 5 | 0.70 ± 0.03 |
| S25 | 3 | 0.63 ± 0.05 | 3 | 0.62 ± 0.03 | 3 | 0.61 ± 0.05 |
| S26 | 4 | 0.67 ± 0.05 | 5 | 0.68 ± 0.02 | 4 | 0.66 ± 0.04 |
| S27 | 4 | 0.59 ± 0.05 | 4 | 0.61 ± 0.04 | 4 | 0.60 ± 0.05 |
| S28 | 5 | 0.67 ± 0.03 | 5 | 0.69 ± 0.02 | 4 | 0.66 ± 0.03 |
| S29 | 4 | 0.78 ± 0.03 | 4 | 0.69 ± 0.02 | 4 | 0.76 ± 0.07 |
| S30 | 4 | 0.65 ± 0.04 | 4 | 0.67 ± 0.03 | 4 | 0.62 ± 0.04 |
| S31 | 4 | 0.67 ± 0.04 | 4 | 0.68 ± 0.02 | 4 | 0.65 ± 0.04 |
| S32 | 4 | 0.66 ± 0.04 | 4 | 0.67 ± 0.03 | 4 | 0.70 ± 0.04 |
| Statistics<br>(mean ± std) | 4 ± 0.69 | 0.68 ± 0.05 | 4 ± 0.76 | 0.67 ± 0.04 | 4 ± 0.69 | 0.67 ± 0.05 |

## Appendix III: Self-Adaptive Threshold Reassignment

In the DEAP database, the emotional ratings of valence and arousal were collected on a continuous scale ranging from 1 to 9, where 1 and 9 indicated the lowest and highest degree of valence or arousal, respectively. Traditionally, a fixed threshold of 5 is directly used to generate binary emotional groups, e.g., low (<5) valence/arousal group and high (>5) valence/arousal group. However, the ranges of the returned self-assessment ratings could be very different from different subjects (as shown in Fig. S3), due to individual-specific experience of emotions (Kemp et al. 2004; Lee et al. 2005). A two-class grouping using a fixed threshold (e.g., 5) could bring out data unbalance and fail to correctly detect neural differences for each subject. Inspired by Yin et al.’s work (Yin et al. 2017), we introduce a self-adaptive threshold reassignment method to group video-stimulated trials into low valence (or arousal) and high valence (or arousal) for each subject. More specifically, the self-adaptive threshold reassignment method for each subject mainly follows 3 steps:

1. ***Individual-level clustering***. A classical k-means clustering is first conducted to partition 40 ratings of one subject into 2 clusters based on two-dimensional valence-arousal space.
2. ***Threshold calculation***. A self-adaptive threshold is then computed as the midpoint of these two cluster centroids.
3. ***Binary emotional grouping***. Based on the computed threshold, 40-trial EEG data collected in the video-stimulated stage are separately divided into low/high valence and low/high arousal groups at the individual level.

An example of the self-adaptive threshold reassignment process on subject 4 is given in Fig. S4. More details about the calculated individual thresholds of valence and arousal for 32 subjects are reported in Table S2, which shows slight variants of the individual thresholds across subjects. Based on the calculated thresholds, 40 trials of EEG data within every single subject can be divided into low- and high-level groups separately on valence and arousal dimensions. A detailed illustration of the self-adaptive grouping results for 32 subjects is presented in Fig. S5.

**Fig. S3.**
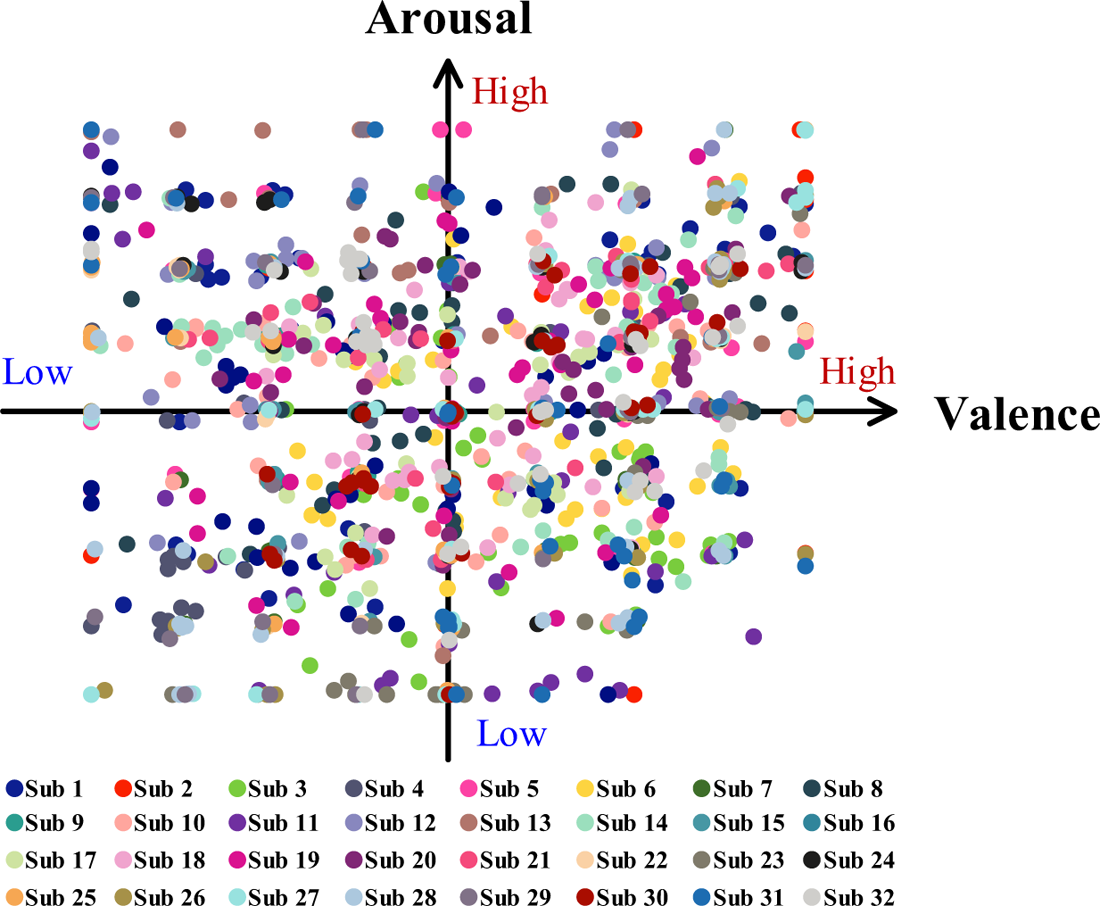
The reported subjective self-assessment ratings on evoked emotions across 32 subjects. The distribution of collected ratings varies from subject to subject, in which a fixed threshold (e.g., 5) would not be suitable for low- and high-level grouping study across 32 subjects.

**Fig. S4.**
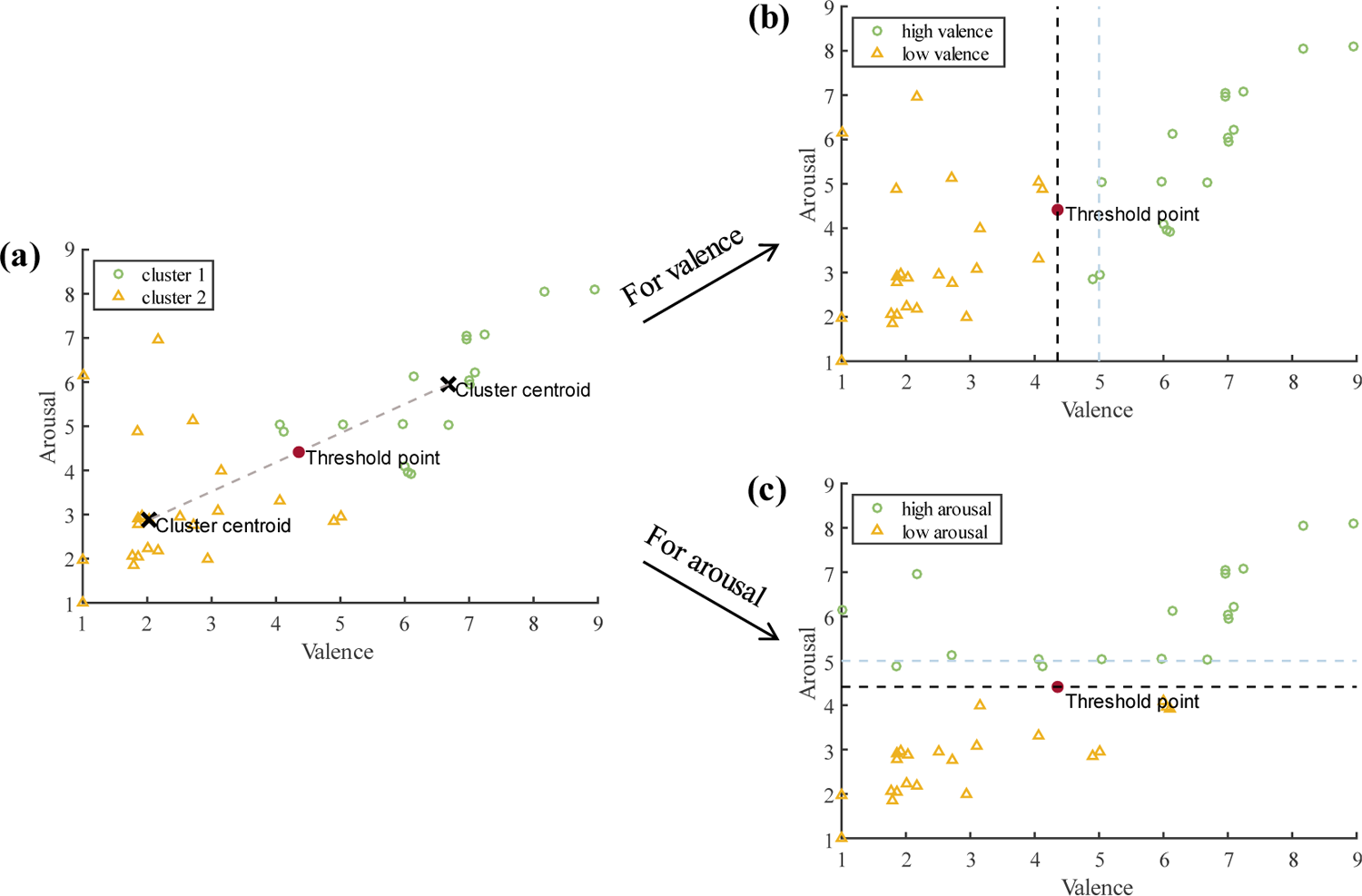
An example of the self-adaptive threshold reassignment method and binary emotional grouping results of subjet 4. (a) The self-adaptive threshold is described as the midpoint of two cluster centroids after a classical k-means clustering. 40-trial EEG recordings are divided into (b) low and high valence groups according to the adaptive threshold on valence dimension, and (c) low and high arousal groups according to the adaptive threshold on arousal dimension.

**Table S2.** The obtained self-adaptive thresholds for subjective ratings on valence and arousal dimensions.

| **Subject ID** | **Valence** | **Arousal** | **Subject ID** | **Valence** | **Arousal** |
| --- | --- | --- | --- | --- | --- |
| **S1** | 5.125 | 6.580 | **S17** | 5.110 | 5.705 |
| **S2** | 6.965 | 4.758 | **S18** | 5.375 | 5.515 |
| **S3** | 5.845 | 3.668 | **S19** | 5.080 | 5.660 |
| **S4** | 4.355 | 4.415 | **S20** | 6.010 | 5.540 |
| **S5** | 5.010 | 4.580 | **S21** | 5.543 | 6.505 |
| **S6** | 6.305 | 4.800 | **S22** | 4.520 | 6.005 |
| **S7** | 5.025 | 5.290 | **S23** | 6.340 | 3.030 |
| **S8** | 6.298 | 5.820 | **S24** | 5.510 | 6.015 |
| **S9** | 5.520 | 5.550 | **S25** | 5.535 | 6.540 |
| **S10** | 5.590 | 5.185 | **S26** | 4.628 | 3.495 |
| **S11** | 5.340 | 4.248 | **S27** | 6.518 | 5.043 |
| **S12** | 4.825 | 6.930 | **S28** | 4.560 | 4.530 |
| **S13** | 5.045 | 7.025 | **S29** | 4.775 | 4.465 |
| **S14** | 4.933 | 6.013 | **S30** | 6.030 | 5.085 |
| **S15** | 6.035 | 4.585 | **S31** | 4.838 | 5.760 |
| **S16** | 4.475 | 4.778 | **S32** | 5.545 | 5.550 |
| **Sample size of low valence** |  | **659** | **Sample size of low arousal** |  | **637** |
| **Sample size of high valence** |  | **621** | **Sample size of high arousal** |  | **643** |
| **Mean** | **self-adaptive** | **threshold** | **for** | **valance** | **ratings** |
| **5.394** |  |  |  |  |  |
| **Mean** | **self-adaptive** | **threshold** | **for** | **arousal** | **ratings** |
| **5.271** |  |  |  |  |  |

**Fig. S5.**
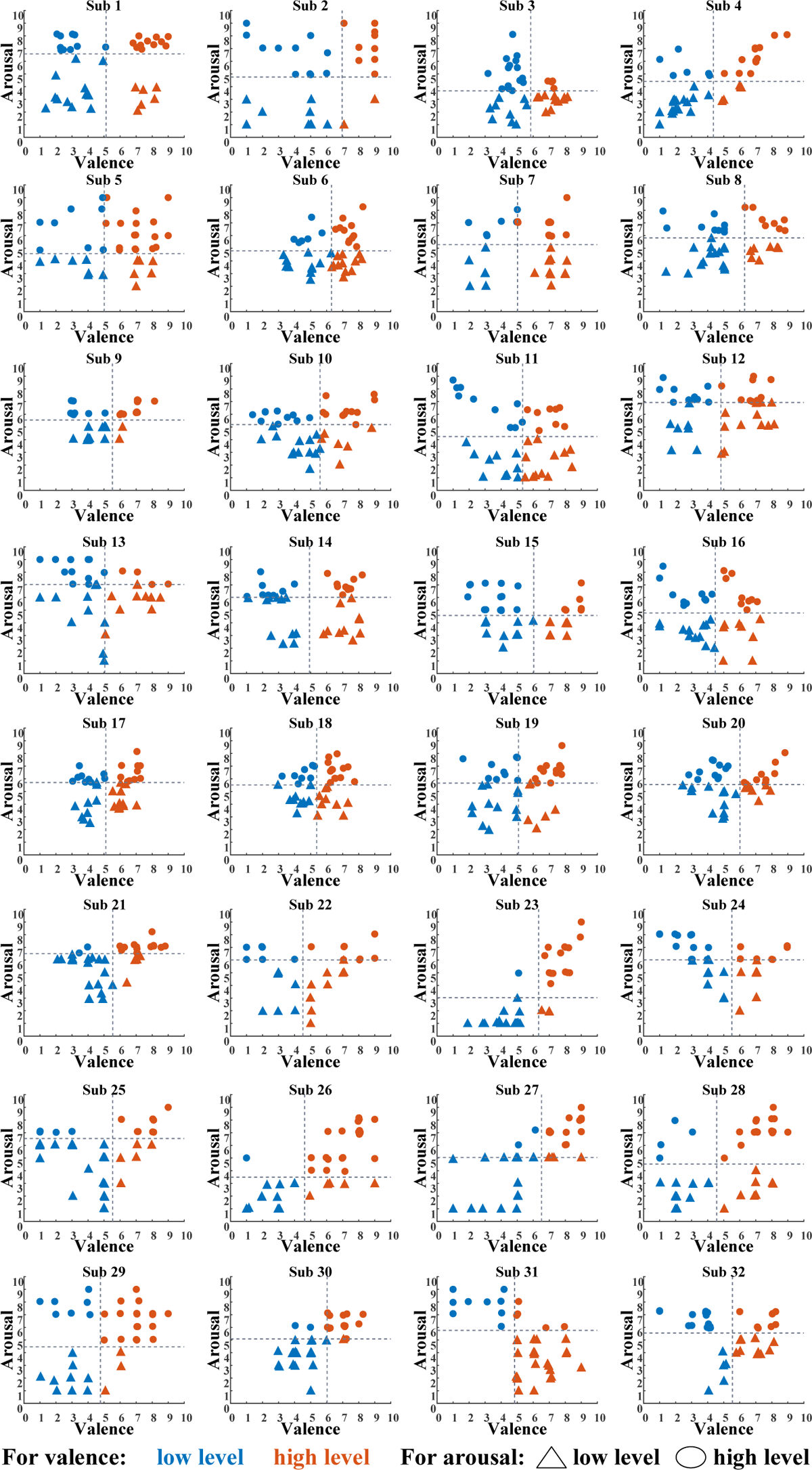
Two-class grouping results on valence and arousal dimensions of 32 subjects using the self-adaptive threshold reassignment method. For better visualization of level differences on valence dimension, the low-level valence rates are marked by blue while the high-level valence rates are marked by orange. For better visualization of level differences on arousal dimension, the low-level arousal rates are marked as a triangle while the high-level rates are marked as circular.

## Appendix IV: Emotion-Evoking Temporal Dynamics in Microstate Sequences

The dynamic changes in MS3 and MS4 coverage of 40 videos across 32 subjects are shown in Fig. S6 and Fig. S7, respectively.

**Fig. S6.**
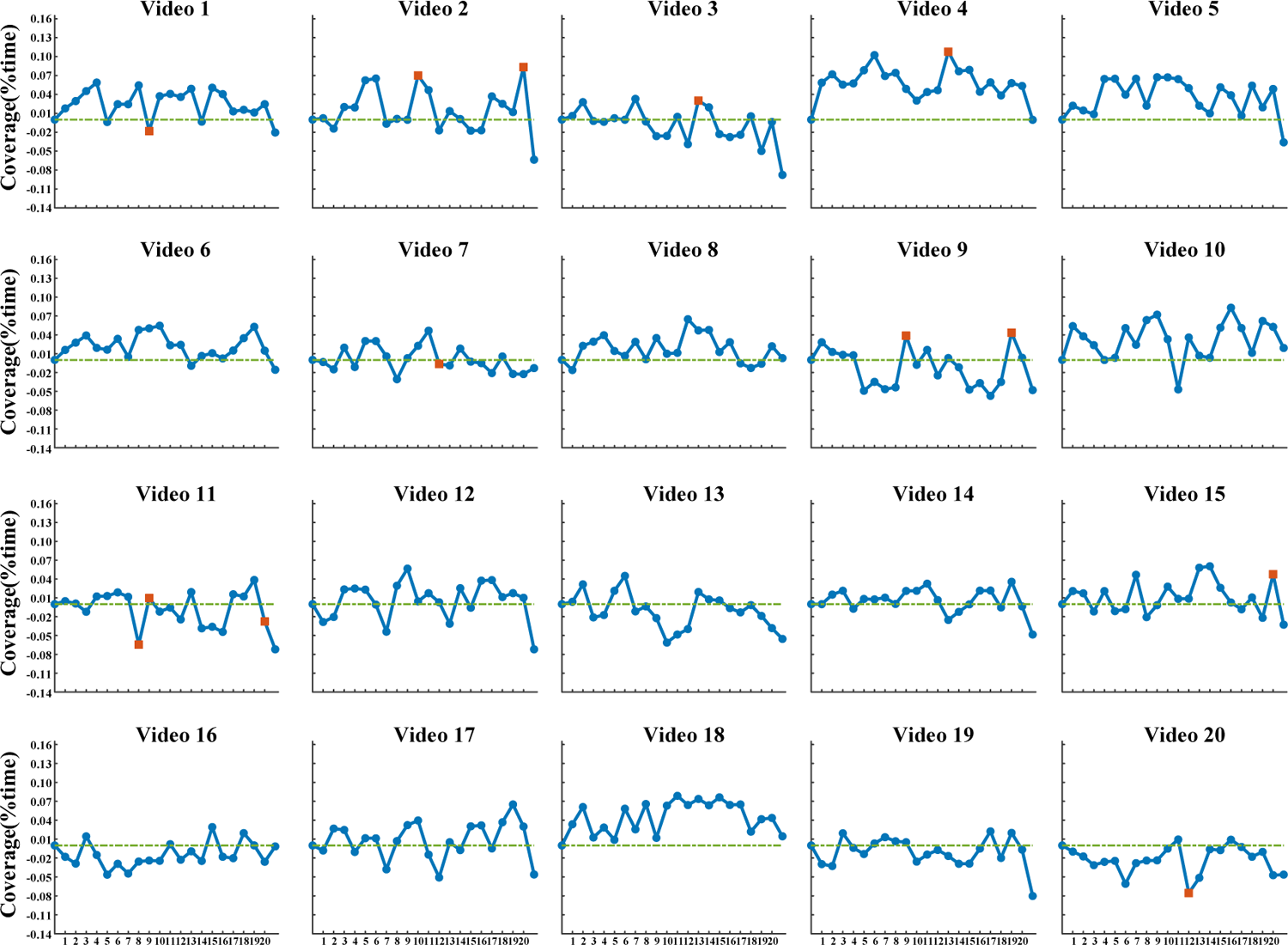
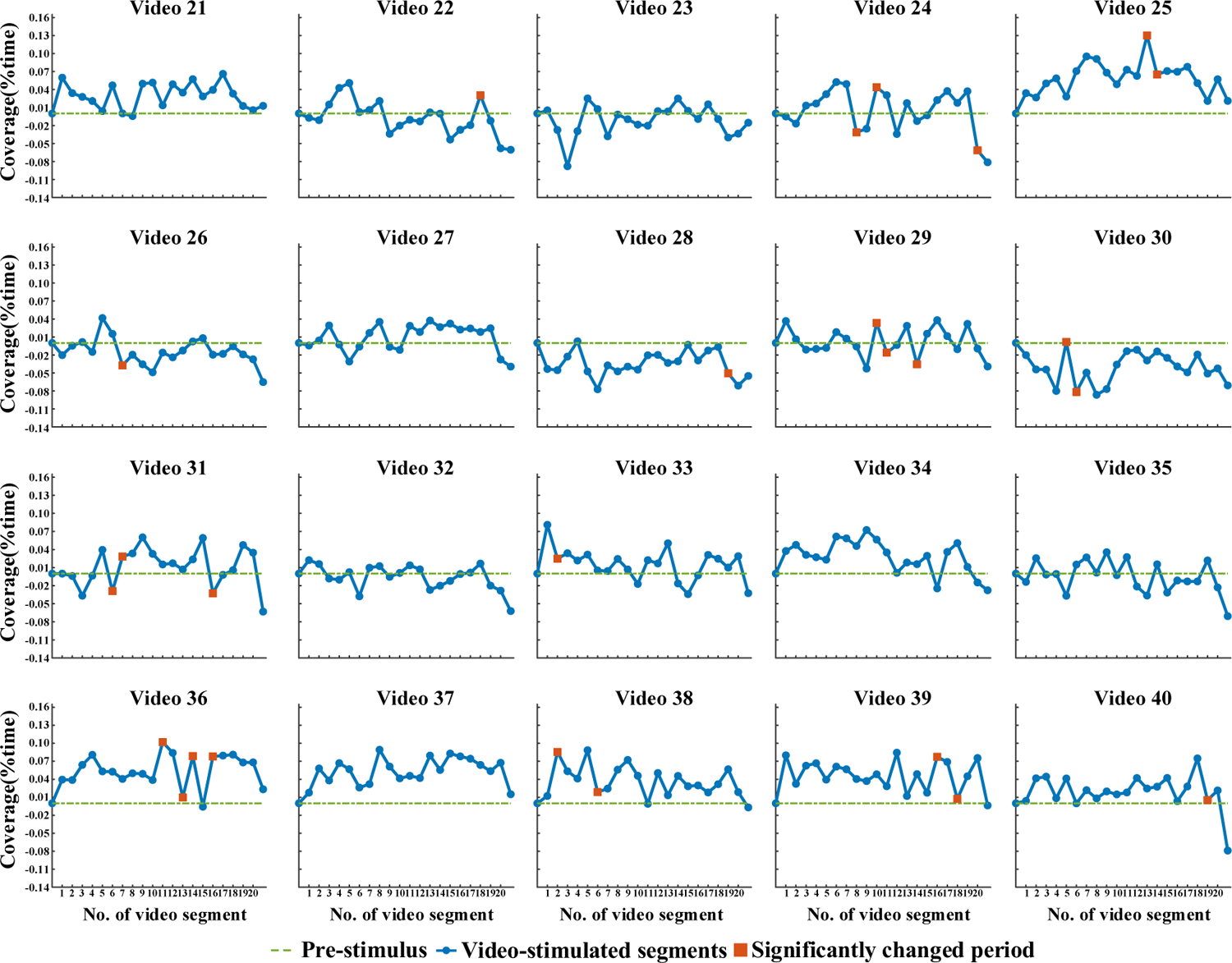
(continued on the next page) The temporal dynamics in MS3 coverage during emotion-evoking under 40 videos. The blue line represents the average MS3 coverage across 32 subjects along with video-triggered emotion-evoking, which shows a general time-varied trend of microstate dynamics evoked by one video. The red square marks the turning points that MS3 coverage extracted from this segment is statistically different from the previous segment across 32 subjects.

**Fig. S7.**
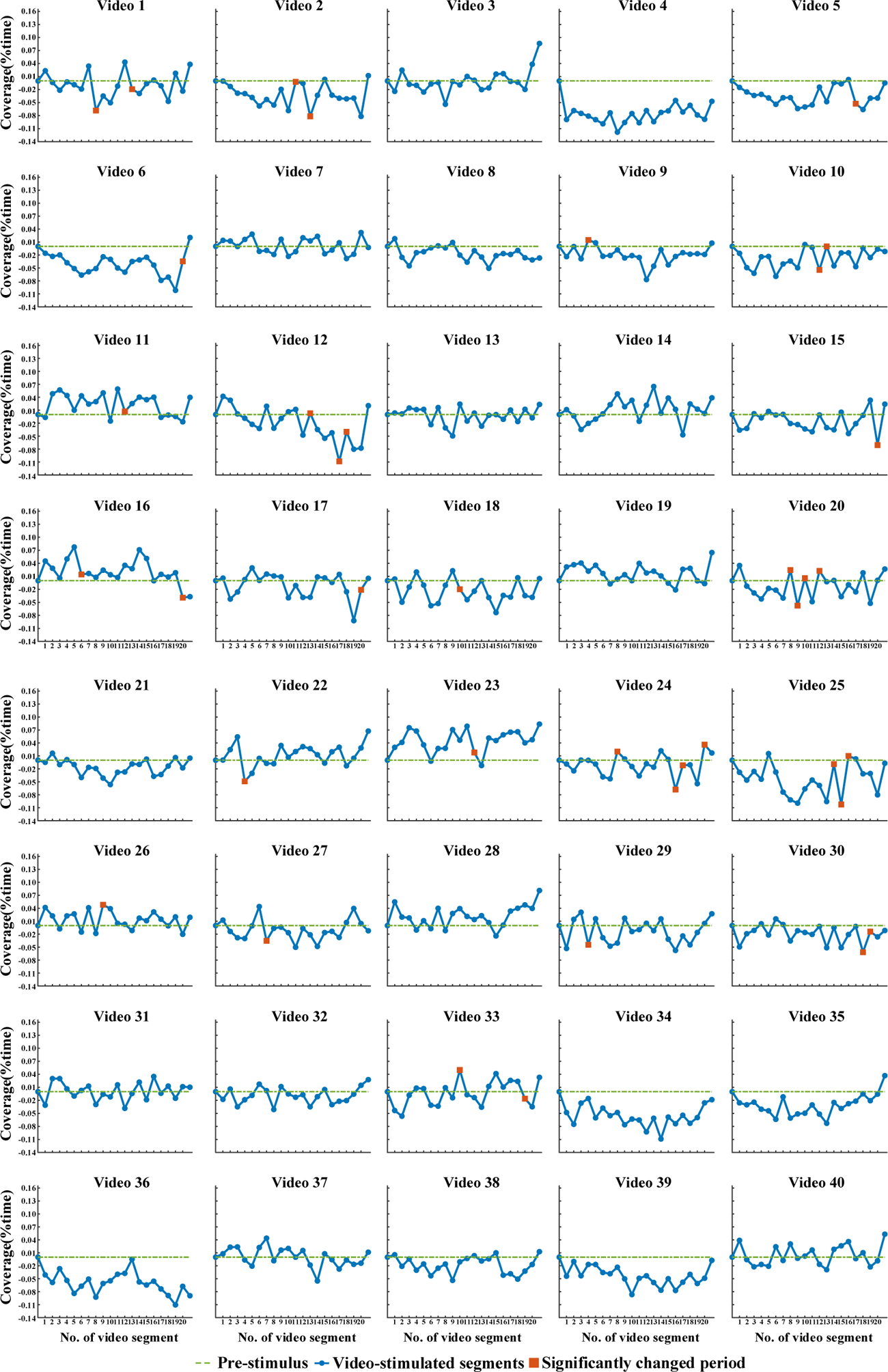
The temporal dynamics in MS4 coverage during emotion-evoking under 40 videos. The blue line represents the average MS4 coverage across 32 subjects along with video-triggered emotion-evoking, which shows a general time-varied trend of microstate dynamics evoked by one video. The red square marks the turning points that MS4 coverage extracted from this segment is statistically different from the previous segment across 32 subjects.

